# Pto evolved from malectin-like RLKs phosphorylates AvrPtoB to promote Prf-mediated immunity in *Solanum pimpinellifolium*

**DOI:** 10.64898/2026.08.23.746126

**Authors:** Lu Liu, Xianyu Zhang, Zhen Gong, Jiancheng Shi, Qingjie Chen, Wei Wu, Jie Ye, Wei Wang, Jun Liu, Ning Xu

## Abstract

Bacterial speck, caused by *Pseudomonas syringae* pv. *tomato* (*Pst*), is a devastating disease of tomato that severely limits global tomato productivity. Understanding the molecular mechanisms underlying *Pst* and tomato is essential for developing disease resistant varieties. Here, we demonstrate that *Pto*, the first disease-resistance gene conferring recognition of a specific pathogen, phosphorylates *Pst* type III effector AvrPtoB at serine 335 site. This post-translational modification triggers the dissociation of the Prf immune complex, enhancing immune signaling and reducing bacterial pathogenicity. Furthermore, evolutionary analyses indicate that Pto-associated proteins originated from malectin-like receptor kinases (MLRs) through loss of the extracellular domain. Crucially, we identified two key amino acid substitutions, Arg158 and Glu258 in Pto, which replace the ancestral lysine residues in MLRs (SpHREK1-1, SpHERK1-2 and SpHERK1-3). These substitutions stabilize Pto by preventing degradation mediated by AvrPtoB’s E3 ubiquitin ligase activity. Our findings reveal a novel mechanism, by which Pto phosphorylates a bacterial effector to trigger enhanced immunity and elucidate the key evolutionary adaptations that have shaped Pto into a stable resistance protein.

**IN A NUTSHELL:** *Background:* The effector AvrPtoB secreted by *Pseudomonas syringae* pv. *tomato* (*Pst*) is recognized by the *Solanum pimpinellifolium* Pto/Prf resistance complex, leading to the dissociation of Pto/Prf resistance complex and activation of the effector-triggered immunity (ETI) response. In *Arabidopsis*, the Ser335 site of AvrPtoB is phosphorylated by the lectin receptor-like kinase LecRK-IX.2, leading to the alleviation of AvrPtoB-mediated suppression of pattern-triggered immunity (PTI). However, in tomato the roles of phosphorylation of AvrPtoB by Pto in the Prf-mediated ETI response remain unclear. In addition, the evolutionary origin of Pto in plants is still poorly understood.

*Question:* How does the phosphorylation of AvrPtoB by Pto activate Prf-mediated ETI response? What is the evolutionary origin of Pto in plants?

*Findings:* Pto phosphorylates AvrPtoB at Ser335 site. AvrPtoB^S335D^ (phosphomimetic type of AvrPtoB) induces programmed cell death (PCD) on RG-PtoR tomato leaves, and exhibits a stronger ability to dissociate the Prf complex compared with AvrPtoB. Moreover, this induction also depends on Pto family protein Pth2. In addition, phylogenetic analysis reveals that malectin-like receptor-like kinase (MLRs) lost their extracellular domain during evolution to give rise to Pto-associated proteins. The tomato MLRs family SpHERK1s show higher homology to Pto than other members and positively regulate resistance to *Pst*. Moreover, Pto Arg158 and Glu258 sites play critical roles in protecting it from AvrPtoB-mediated degradation.

*Next steps:* The intrinsic mechanism by which phosphorylation at the Ser335 site of AvrPtoB activates the Prf-mediated ETI response deserves in-depth exploration. Additionally, the effect of SpHERK1-mediated phosphorylation of AvrPtoB at Ser335 on the Prf-mediated ETI response deserves further exploration.

## Introduction

*Pseudomonas syringae* pv. tomato (*Pst*) is one of the most extensively studied plant pathogens, which serves as a model system for investigating bacterial pathogenicity and molecular mechanisms underlying plant–microbe interactions (Xin and He, 2013). The interaction between *Pst* and tomato leads to bacterial speck disease, which can cause severe yield losses from 10%-30%, with the most severe cases resulting in up to 100% yield loss (Wang et al., 2018). Plants have evolved two major strategies to mount an immune response against pathogens. The first layer is pattern-triggered immunity (PTI), which is activated by conserved pathogen-, damage-, microbe-, or herbivore-associated molecular patterns (PAMPs/DAMPs/MAMPs/HAMPs) via recognition by cell surface-localized pattern recognition receptors (PRRs). The second layer is effector-triggered immunity (ETI), initiated by the recognition of pathogen effectors via intracellular nucleotide-binding leucine-rich repeat receptors (NLRs) (Jones and Dangl, 2006). PTI is faster than ETI, while ETI is highly robust and typically results in stronger immunity activation. Rather than two independent systems, recent studies have demonstrated the mutual reinforcement of PTI and ETI (Ngou et al., 2021; Yuan et al., 2021).

NLRs can be classified into three classes according to their N-terminal domains: coiled-coil (CC) NLRs (CNLs), Toll/Interleukin-1 receptor/Resistance (TIR) protein NLRs (TNLs), and RPW8-like CC domain (RPW8) NLRs (RNLs). To date, the only NLR protein that confers resistance to bacterial speck in tomato is the CNL protein Prf (Salmeron et al., 1996), which can form immune complexes with the intracellular serine/threonine kinase protein Pto (Saur et al., 2015). Pto directly interacts with two type III effectors from *Pst*, AvrPto and AvrPtoB (Kim et al., 2002). Recognition of AvrPto/AvrPtoB by the Prf/Pto complex triggers ETI responses, including mitogen-activated protein kinase (MAPK) signaling cascades and programmed cell death (PCD) (Abramovitch et al., 2003; del Pozo et al., 2004; Pedley and Martin, 2004). Recent advances have elucidated the mechanism underlying the interaction between helper NLRs (NRC2, NRC4) and the Prf-/Pto complex during ETI (Liu et al., 2024a; Madhuprakash et al., 2024). Following effector (AvrPto/AvrPtoB) recognition, the complex dissociates into two distinct ETI signaling branches: the MAPK-branch and the NRC-branch. The NRC-branch is essential for most PCD formation and for full restriction of pathogen growth (Sheikh et al., 2023). In the *nrc2*/*nrc3* tomato mutant, Prf/Pto-mediated ETI is abolished in response to *Pst* delivering AvrPto and AvrPtoB (Zhang et al., 2024).

AvrPtoB is a 553-amino-acid protein containing a functional E3 ligase domain at its C terminus. Recent studies have shown that AvrPtoB targets multiple PRRs, helper NLRs, and components of hormone signaling pathways to promote infection (Göhre et al., 2008; Gimenez-Ibanez et al., 2009; Chen et al., 2017; Xu et al., 2020; Liu et al., 2022; Wang et al., 2023). These findings underscore the role of AvrPtoB as a major virulence effector in *Pst*. To counter this pathogenicity, plants have evolved mechanisms to attenuate the virulence of AvrPtoB. For example, in tomato, Pto phosphorylates AvrPtoB at threonine 450 site to inhibit the E3 ligase ability (Ntoukakis et al., 2009). In *Arabidopsis*, L-type lectin receptor-like kinase (RLK) LecRK-IX.2 interacts with and phosphorylates AvrPtoB at serine 335 to prevent AvrPtoB self-association and to mitigate E3 ligase-mediated degradation (Xu et al., 2020).

The bacterial speck disease-resistant protein Pto was originally cloned from a wild relative of tomato *Solanum pimpinellifolium* (Martin et al., 1993). Subsequent studies identified gene families with high homology to Pto in other Solanaceous species. In tomato, all accessions from 12 wild relatives possess the Pto gene family, and the orthologs across species exhibit high sequence similarity (Rose et al., 2005). Natural variations in Pto have been observed; for instance, *Solanum chmielewskii* (Schm) accessions are susceptible to *Pst* expressing AvrPto but resistant to strains expressing AvrPtoB. This phenotype is attributed to a single histidine-to-aspartate substitution at position 193 in Pto, which is sufficient to confer recognition of AvrPto in tomato (Kraus et al., 2016).

In pepper, 25 Pto-like protein kinases (PLPKs) containing a highly conserved serine-threonine kinase (STK) domain have been identified (Venkatesh et al., 2016). In *Nicotiana. benthamiana*, *NbPth1* to *NbPth3* were amplified using primers based on *Pto* sequences (Mucyn et al., 2006), and silencing of *NbPth1* abolishes the AvrPto/Pto-dependent HR (Gutierrez et al., 2010). Interestingly, primers designed for *Pto* could also amplify candidates of *Catharanthus roseus* receptor-like kinase 1-like proteins (CrRLK1Ls) from *Platanus*. Previous studies revealed that the Pto group is more closely related to the CrRLK1L subfamily than to other RLK subfamilies, suggesting a possible evolutionary origin of Pto from CrRLK1L (Pilotti et al., 2014). However, the precise evolutionary mechanism of Pto remains unclear.

CrRLK1Ls belong to the malectin-like receptor kinase (MLR) family and are widespread in monocots and dicots, participating in diverse plant physiological processes (Franck et al., 2018). All CrRLK1L members share a conserved structure comprising one or two extracellular malectin-like domain (MLD), a transmembrane domain (TMD), and an intracellular serine/threonine kinase domain. The well-characterized member is FERONIA (FER) in *Arabidopsis*, which serves as a receptor for rapid alkalinization factor (RALF) peptides (Haruta et al., 2014). FER regulates numerous processes including pollen hydration (Liu et al., 2021), pavement cell morphogenesis (Lin et al., 2022), abiotic stress responses (Liu et al., 2024b), mRNA translation (Xu et al., 2024), and plant–pathogen interactions (Stegmann et al., 2017;

Zhang et al., 2020). In tomato, SlFERL recognizes the virulence factor BcPG1 from *Botrytis cinerea* via its extracellular domain and activates MAPK immune signaling by phosphorylating SlMAP3K18 (Ji et al., 2023). Nevertheless, the functions of other malectin-like kinases in tomato remain largely unexplored.

In this study, we provide evidence that Pto directly phosphorylates AvrPtoB at serine 335 site. This specific phosphorylation potently disrupts the Pto/Prf immune complex, leading to its dissociation and triggering an immunity response. Furthermore, through phylogenetic and structural analyses, we demonstrate that Pto and its homologs evolved from ancestral malectin-like receptor kinases (MLRs) via loss of the extracellular malectin-like domain. Notably, we identified two critical amino acid substitutions in Pto, Arg158 and Glu258, which replace lysine residues conserved in ancestral MLRs such as SpHERK1s. These substitutions are essential for Pto protein stability, as they prevent targeted degradation mediated by the E3 ubiquitin ligase activity of AvrPtoB. Collectively, our study uncovers a novel mechanism in which host phosphorylation of a bacterial effector boosts immunity, and delineates the key evolutionary innovations that stabilized Pto, enabling its specialized function as a disease resistance protein.

## Results

### Pto phosphorylates serine at 335 of AvrPtoB

In our previous study, we demonstrated that *Arabidopsis thaliana* LecRK-IX.2 phosphorylates *Pst* effector AvrPtoB at Ser335 site to attenuate its virulence. Structural modeling revealed that the kinase domain (KD) of LecRK-IX.2 shares similarity with that of Pto (Xu et al., 2020). Given that Pto was previously shown to phosphorylate AvrPtoB at Thr450 (Ntoukakis et al., 2009), we performed *in vitro* kinase assays to test whether Pto can also phosphorylate AvrPtoB at Ser335. We cloned *Pto* from the wild tomato relative *Solanum pimpinellifolium* and the cultivated species *S. lycopersicum* for kinase assays. Using site mutants of AvrPtoB^S335A^ and AvrPtoB^T450D^ as negative control, we found that both Pto variants phosphorylated AvrPtoB and AvrPtoB^T450D^ at Ser335, as detected by anti-pS335 antibody (Fig. 1, A and B). Fen, a homolog of Pto with 87% amino acid similarity (Martin et al., 1994), also phosphorylated AvrPtoB at Ser335 *in vitro* (Fig. 1C). Consistent with earlier reports, Fen exhibited weaker general kinase activity toward AvrPtoB by α-pSer/Thr testing (Ntoukakis et al., 2009). Still it showed the same phosphorylation efficiency at Ser335 as SlPto (Fig. 1D). Kinase-dead mutants of Pto, SlPto, and Fen were used as negative control (Supplementary Fig. S1, A to C). Next we co-expressed *35S:AvrPtoB-HA* with *35S:Pto-T7* or *35S:Pto^D164N^*-*T7* in *N. benthamiana*. The result showed that Pto could phosphorylate AvrPtoB at Ser335 *in vivo*. Pto^D164N^ (Kinase-dead mutant) was used as a negative control (Supplemental Fig. S1D).

**Figure 1.**
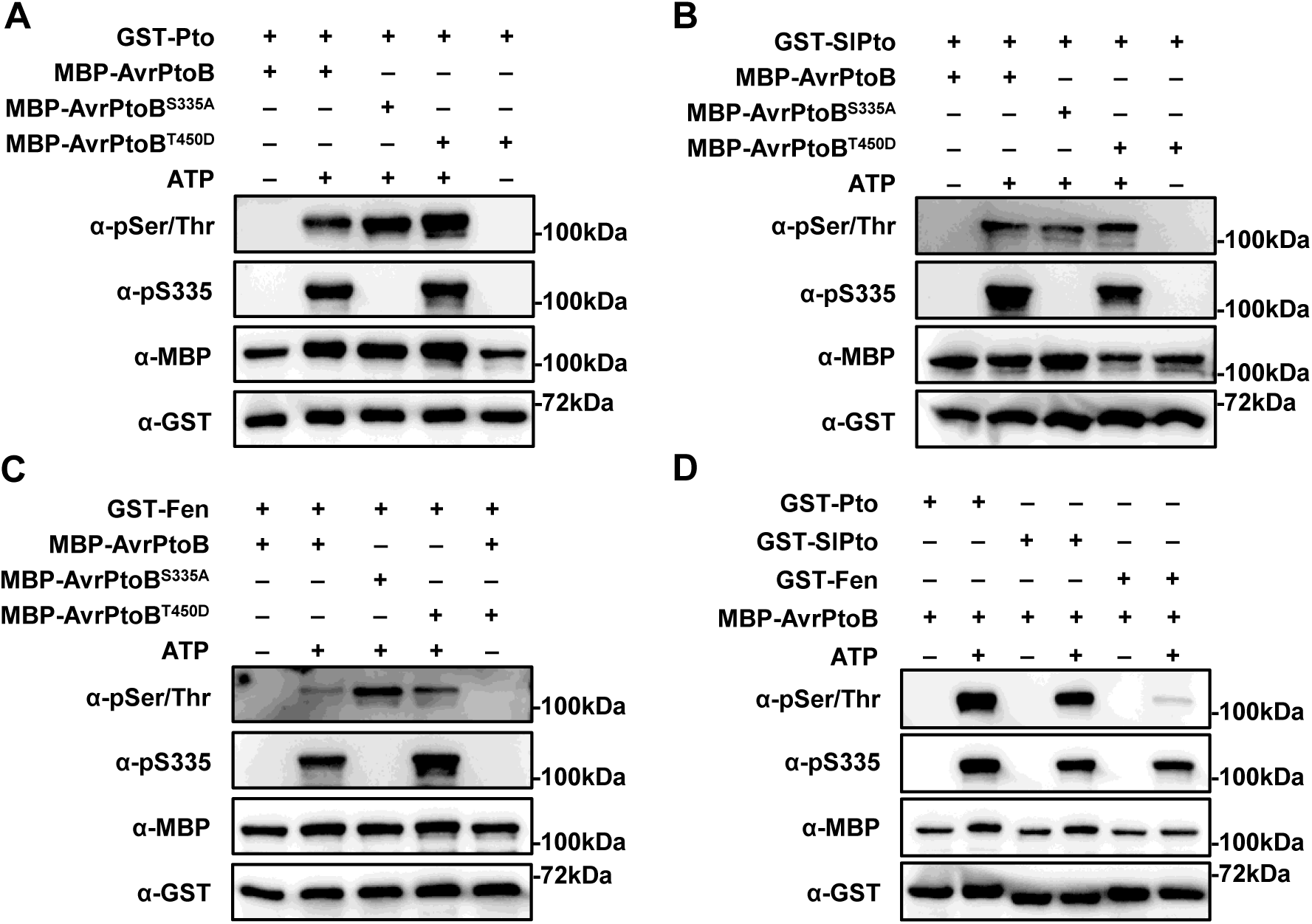
Pto Phosphorylates AvrPtoB at Ser335. **(A–D)** Pto and Fen phosphorylate AvrPtoB at Ser335 *in vitro* kinase assay. Recombinant GST-Pto (from *S. pimpinellifolium*), GST-SlPto (from *S. lycopersicum*), GST-Fen (from *S. pimpinellifolium*), MBP-AvrPtoB and site-mutated variants AvrPtoB^S335A^ and AvrPtoB^T450D^ were subjected to kinase assay. Protein phosphorylation was identified by immunoblot using anti-phosphoserine/threonine antibody (α-pSer/Thr). The phosphorylated AvrPtoB was detected by anti-pS335 antibody (α-pS335).

To investigate whether Ptos from other species have such phosphorylation activity, we cloned *Pto* genes from 1 cultivated tomato (*Solanum lycopersicum* MoneyMaker) and another 10 wild tomatoes (*S. lycopersicoides*, *S. habrochaites*, *S. pennellii*, *S. chilense*, *S. peruvianum*, *S. corneliomulleri*, *S. neorickii*, *S. chmielewskii*, *S*. *pimpinellifolium*, and *S. galapagense*) (Rose et al., 2005; Kraus et al., 2016; Li et al., 2023). Sequences alignment revealed 80.56% to 96.86% identity among these Pto homologs. Notably, all Ptos contained a conserved kinase domain, an activation domain, and P+1 loop, a vital site that mediates the Pto-AvrPto (B) interaction. The ATP binding site (Lys-69 of Pto) was also highly conserved in different Pto proteins (Supplementary Fig. S2). Kinase assay confirmed that all Pto homologs phosphorylated AvrPtoB at Ser335 (Supplementary Fig. S3). Taken together, these results indicate that phosphorylation of AvrPtoB at Ser335 is a conserved function among diverse Pto variants.

### Phosphorylation of AvrPtoB at Ser335 site reduces the pathogenicity of *Pst*

As a well-studied E3 ligase, the N-terminus of AvrPtoB can cause PCD, while the C-terminus can function as an E3 ligase to inhibit PCD (anti-PCD). In RG-PtoR, which harbors the resistance proteins Pto and Prf, Pto can block the E3 function of AvrPtoB and promote PCD. In RG-pto11 (*pto* mutant line), which lacks PtoR, the N-terminus induced PCD can be blocked by its C-terminus, so there is no PCD in RG-pto11 line. Mutation of conserved residue of AvrPtoB Phe479, which is involved in the binding of E2 ubiquitin–conjugating enzymes, can abolish its anti-PCD activity and trigger Prf-dependent PCD in RG-pto11 (Abramovitch et al., 2003; Abramovitch and Martin, 2005; Abramovitch et al., 2006; Janjusevic et al., 2006; Ntoukakis et al., 2009). To explore the biological relevance of Pto-mediated phosphorylation of AvrPtoB at Ser335, we infiltrated *Agrobacterium* GV2260 strains expressing AvrPtoB, AvrPtoB^S335D^ (phosphomimetic type of AvrPtoB), AvrPtoB^S335A^ or AvrPtoB^F479A^ in RG-PtoR, RG-pto11 and RG-prf3 (*prf* mutant line) tomato lines, respectively. After three days post inoculation (dpi), all four AvrPtoB varieties elicited PCD in RG-PtoR lines, while only AvrPtoB^S335D^ and AvrPtoB^F479A^ induced PCD in RG-pto11 lines. None caused PCD in RG-prf3 plants (Fig. 2A), indicating that AvrPtoB^S335D^ triggered PCD depends on Prf.

**Figure 2.**
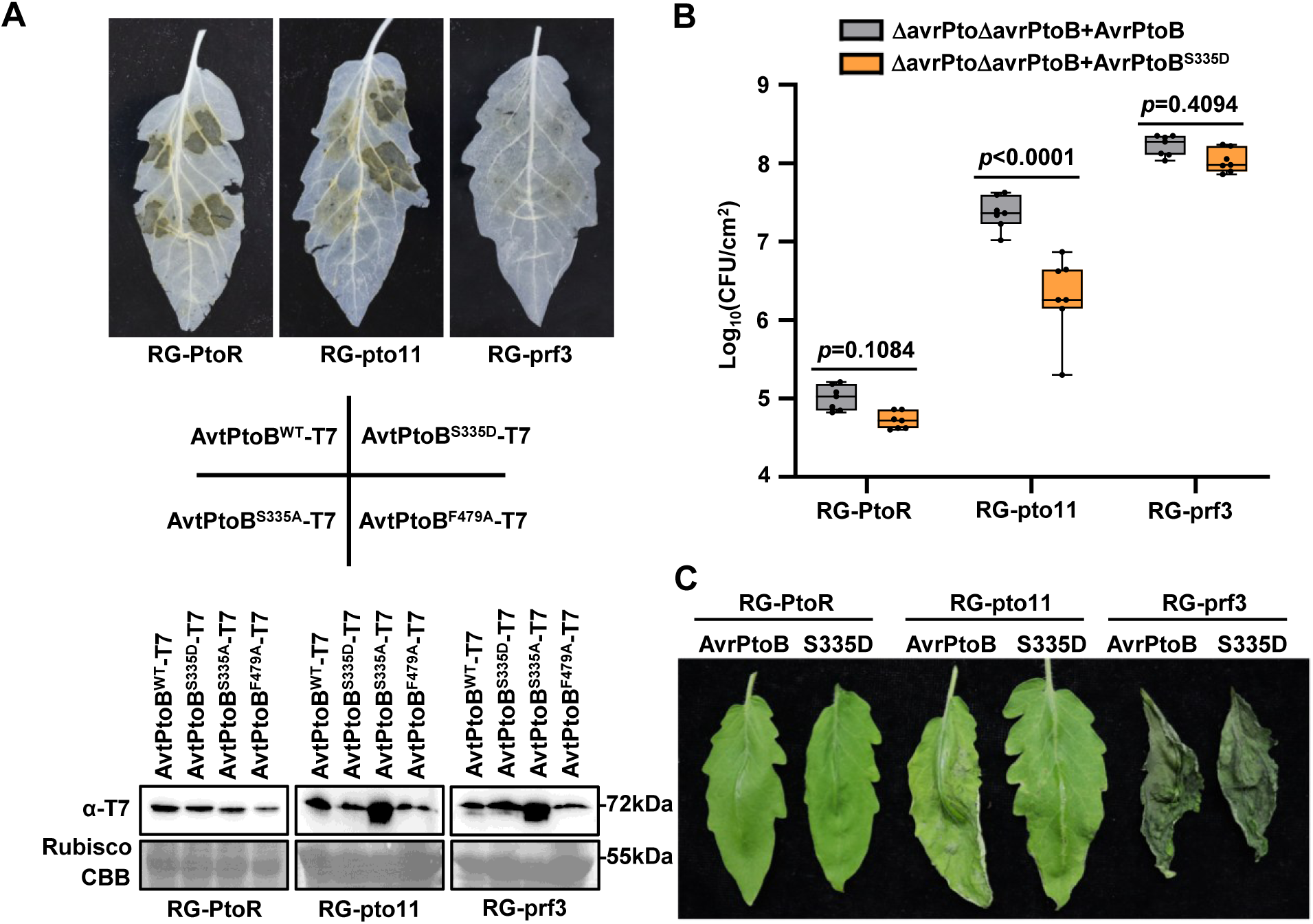
Phosphorylation of AvrPtoB at Ser335 by Pto reduces the pathogenicity of *Pst* DC3000. **(A)** AvrPtoB^S335D^ triggers enhanced cell death in tomato plants*. Agrobacterium* GV2260 harboring *35S:AvrPtoB-T7*, site-mutated variants *AvrPtoB^S335D^-T7*, *AvrPtoB^S335A^-T7* and *AvtPtoB^F479A^-T7* were infiltrated into RG-PtoR, RG-pto11 and RG-prf3 tomato leaves. Ethanol was applied for decolonization after 72 hpi. Down panel showing the protein expression levels. CBB-stained Rubisco was used as loading control. The experiment was repeated at least three times with similar results. **(B)** Phosphorylation mimic of S335 impairs AvrPtoB virulence in RG-pto11. Four-week-old tomato plants were inoculated with *Pst* DC3000 ΔavrPtoΔavrPtoB expressing AvrPtoB and AvrPtoB^S335D^ at the concentration of 5×10^4^ CFU/mL. The plants were subjected to growth curve analysis at 3 dpi. Error bars represent means ± SD (two-way ANOVA, n = 7). *p-*values were marked on the bar. **(C)** Disease phenotype of tomato plants. Four-week-old tomato plants were sprayed with *Pst* DC3000 ΔavrPto*Δ*avrPtoB expressing AvrPtoB and AvrPtoB^S335D^ at the concentration of 2×10^8^ CFU/mL. Pictures were taken at 3 dpi.

Given the redundant avirulence activities of AvrPto and AvrPtoB in Pto-expressing tomatoes (Lin and Martin, 2005), we generated *Pst Δ*avrPtoΔavrPtoB double mutant to complement AvrPtoB and AvrPtoB^S335D^ for pathogenicity assays. In RG-PtoR, there was no significant difference in pathogenicity between *Pst* ΔavrPtoΔavrPtoB double mutant expressing AvrPtoB and AvrPtoB^S335D^. In RG-pto11, the pathogenicity of *Pst* ΔavrPtoΔavrPtoB double mutant expressing AvrPtoB^S335D^ was significantly reduced compared to AvrPtoB. In RG-prf3, no significant difference in pathogenicity was observed between *Pst* ΔavrPtoΔavrPtoB expressing AvrPtoB and AvrPtoB^S335D^ (Fig. 2, B and C), suggesting that AvrPtoB^S335D^ induced PCD and disease resistance are not entirely dependent on Pto and entirely dependent on Prf.

### Phosphorylation of AvrPtoB at Ser335 promotes dissociation of Prf/Pto resistance complex

Previous studies have verified that Prf/Pto resistance complex is oligomeric and contains Pto or one of several Pto kinase family members, and at least two Prf molecules (Gutierrez et al., 2010). Following AvrPto/AvrPtoB recognition, Pto/Prf complex dissociates and triggers downstream immune responses, including activation of MAPK cascade, PCD, and the restriction of pathogen growth (Sheikh et al., 2023) (Supplementary Fig. S4A). Prf consists of an N-terminal domain (Nterm) and an SD-CC-NB-NRC-LRR structure (SCNL) (Fig. 3A). Co-expression of Prf’s N-term and SCNL in *N. benthamiana* reconstitutes the fully functional Prf. The N-term of Prf specifies homotypic interactions independently of Pto, but the C-term can’t (Gutierrez et al., 2010). AvrPtoB can promote the dissociation of Prf’s N-term in Split-Luciferase Complementation Assay (LCA) (Sheikh et al., 2023).

**Figure 3.**
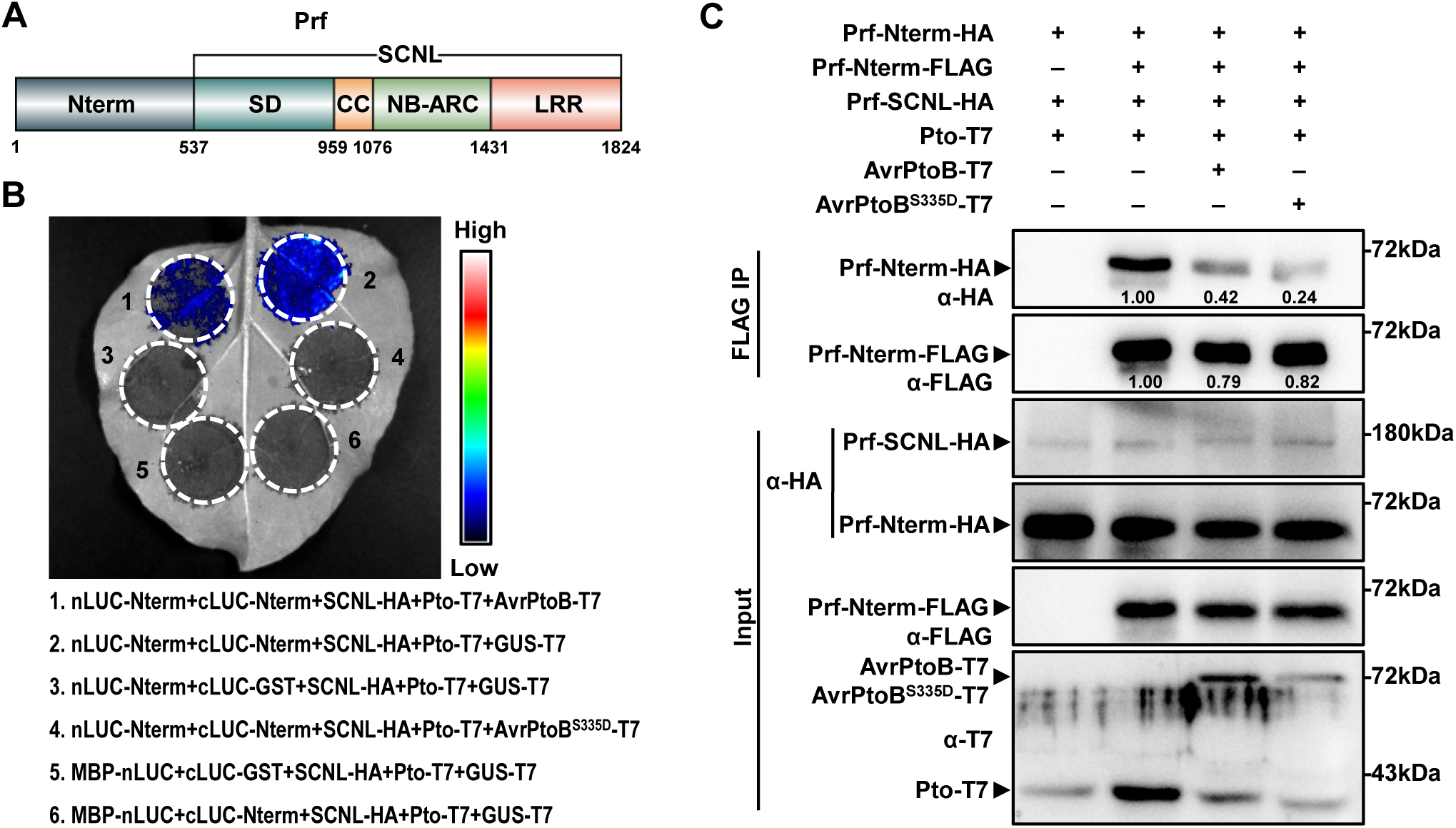
Phosphorylation of AvrPtoB at Ser335 promotes the dissociation of Prf-Prf complex. **(A)** Schematic diagram of Prf. Nterm: N-terminal domain; SD: Solanaceous domain; CC: coiled-coil domain; NB-ARC: Nucleotide-Binding adaptor shared by APAF-1, R proteins, and CED-4; LRR: leucine-rich repeat domain; SCNL: SD-CC-NB-ARC-LRR structure. Numbers indicated the amino acids of Prf protein sequence. **(B)** AvrPtoB^S335D^ promotes dissociation of Prf complex in *vivo* by split-luciferase complementation assays. *35S:nLUC-Prf-Nterm* (Prf N-terminal sequence), *35S:cLUC-Prf-SCNL* (Prf C-terminal sequence), *35S:SCNL-HA*, *35S:Pto-T7* and *35S:AvrPtoB-T7* or *35S:AvrPtoBS^335D^-T7* were transiently expressed in *N. benthamiana* leaves. The chemiluminescence images were obtained by applying 0.5 mM luciferin at 36 hpi (before necrosis occurred). **(C)** AvrPtoB^S335D^ promotes dissociation of Prf complex *in vivo* by Co-IP assay. *35S:Prf-Nterm*, *35S:Prf-SCNL*, *35S:Pto* and *35S:AvrPtoB* or *35S:AvrPtoBS^335D^* were transiently expressed in *N. benthamiana* leaves. Extracted proteins were immunoprecipitated 36 hpi (before necrosis occurred) using anti-FLAG beads. The Co-IP proteins were detected by anti-HA antibody. The band intensity was calculated by Image J. The smaller value represents the weaker interaction between proteins. All experiments were repeated three times with similar results.

To compare the effect of AvrPtoB and AvrPtoB^S335D^ on the dissociation of Pto/Prf complex, we conducted LCA and Co-immunoprecipitations (Co-IP). The LCA result showed that compared with AvrPtoB, AvrPtoB^S335D^ had stronger ability to dissociate the interaction between Prf/Pto resistance complex (Fig. 3B). For Co-IP assay, consistent with the previous research, the interaction between two Prf Nterms was weakened when AvrPtoB was recognized by Pto (Sheikh et al., 2023) (Fig. 3C). Furthermore, AvrPtoB^S335D^ promoted more dissociation of Prf-Prf complex than AvrPtoB (Fig. 3C). Therefore, we hypothesized the phosphorylation at AvrPtoB Ser335 is critical for Prf complex dissociation and PCD induction in tomato.

### Phosphorylation of AvrPtoB at Ser335 mediates Prf complex dissociation via Pth2 in RG-pto11 plants

Although AvrPtoB-induced dissociation of the Prf complex is typically Pto-dependent (Gutierrez et al., 2010), we found that AvrPtoB^S335D^ still elicited PCD in RG-pto11 (Fig. 2A), implying involvement of additional components. Previous studies showed that AvrPtoB^F479A^ or AvrPtoB^1–387^, both deficient in E3 ligase activity, trigger PCD in a Fen-dependent manner. AvrPtoB promotes Fen degradation via ubiquitin-proteasome pathways (Janjusevic et al., 2006; Rosebrock et al., 2007). To investigate whether AvrPtoB^S335D^ induced PCD in RG-pto11 was related to Fen/Prf immune pathway, we first tested the E3 ligase activity of AvrPtoB^S335D^ to Fen. *In vitro* ubiquitination assays indicated that AvrPtoB^S335D^ could ubiquitinate Fen as AvrPtoB (Supplementary Fig. S4, B and C). Pto was used as a negative control. Further experiment demonstrated that AvrPtoB^S335D^ can degrade Fen in planta via 26S proteasome *in vivo* (Supplementary Fig. S4D). To check whether AvrPtoB or AvrPtoB^S335D^ induced dissociation of Prf complex dependent on Fen, we conducted Co-IP assay. Unexpectedly, Fen was not required for AvrPtoB or AvrPtoB^S335D^ induced dissociation of Prf complex (Supplementary Fig. S4E).

Pto is one member of a multigene family, besides Fen, the Pto family contains four other members: Pth2 (PtoD), Pth3 (PtoC), Pth4 (PtoA) and Pth5. Among them, Pth5 is considered to be a pseudogene (Chang et al., 2002; Rosebrock et al., 2007). We hypothesized that other members may participate in this signaling pathway, so we first selected Pth2 for further study. Consistent with the previous study, Pth2 could not be degraded and ubiquitinated by AvrPtoB (Rosebrock et al., 2007) (Fig. 4A; Supplementary Fig. S5A). Fen was use as a positive control. Although previous study showed that Pth2 lacks self-phosphorylation activity *in vitro* (Chang et al., 2002), our kinase assay exhibited that Pth2 had weaker self-phosphorylation activity and phosphorylation ability toward AvrPtoB than Pto (Fig. 4B; Supplementary Fig. S5B). Co-IP assay verified that Pth2 interacted with Prf Nterm, AvrPtoB and AvrPtoB^S335D^ (Fig. 4, C and D). Next we checked whether AvrPtoB^S335D^ dissociated Prf complex via Pth2. The results showed that AvrPtoB^S335D^ had stronger ability to dissociate the interaction between Prf/Pth2 resistance complex (Fig. 4E). Moreover, we used virus induced gene silence (VIGS) technology to knock-down the expression of *Pth2* in RG-pto11 and found that AvrPtoB^S335D^-induced PCD was significantly reduced in *Pth2*-silenced tomato compared with control tomato (Fig. 4F; Supplementary Fig. S5C). Next, we performed the bacterial growth experiment in *TRV:EV* and *TRV:Pth2* in RG-Pto11. The result showed that the recognition of AvrPtoB^S335D^ in RG-pto11 is Pth2-dependent (Supplementary Fig. S5D). We also checked whether Pth3 and Pth4 had the same function like Pth2. Our Co-IP assay showed that AvrPtoB^S335D^ could not dissociated Prf complex through Pth3 compared with Pth2 (Supplementary Fig. S5E). PCR could not amplify *Pth4* from RG-PtoR. Taken together, these results demonstrated that Pth2 participated in AvrPtoB^S335D^-induced PCD in RG-pto11 tomato plants.

**Figure 4.**
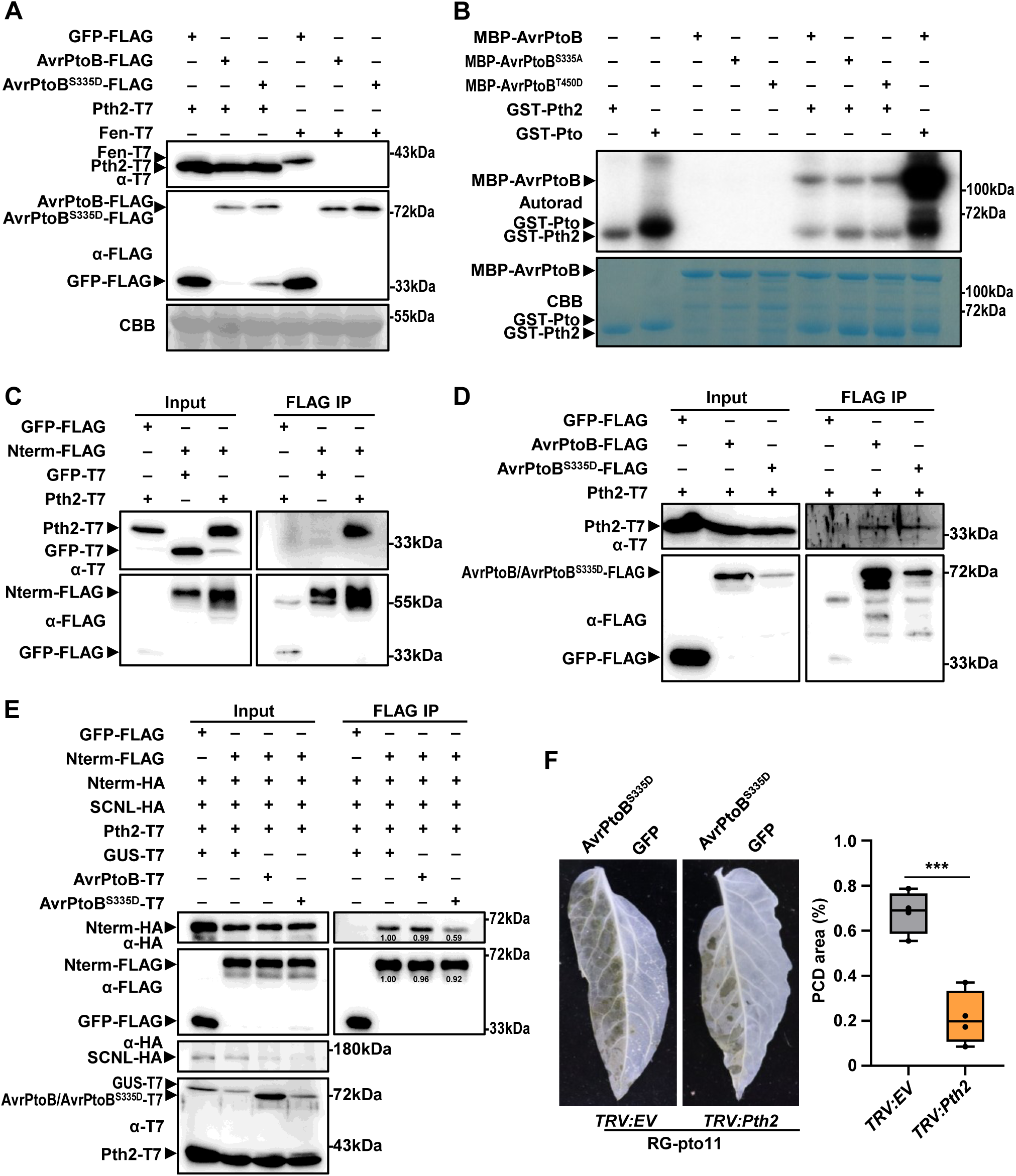
The dissociation of Prf-Prf complex induced by AvrPtoB^S335D^ depends on Pth2. **(A)** AvrPtoB can’t degrade Pth2 *in vivo*. *35S:Pth2-T7*, *35S:AvrPtoB* or *35S:AvrPtoB^S335D^*were transiently expressed in *N. benthamiana* leaves. The proteins were examined by immunoblotting with respective antibodies. Fen was used as positive control. **(B)** Pth2 can phosphorylate AvrPtoB *in vitro* kinase assay. Recombinant MBP-AvrPtoB, MBP-AvrPtoB^S335A^, MBP-AvrPtoB^T450D^, GST-Pth2, and GST-Pto were subjected to kinase assay. Protein phosphorylation was identified by autoradiography. Protein abundance was stained by CBB in SDS–PAGE gel. **(C)** Pth2 interacts with Prf-Nterm *in vivo* by Co-IP assay. *35S:Pth2-T7*, *35S:GFP-T7*, *35S:Prf-Nterm*-*FLAG* or *35S:GFP-FLAG* were transiently expressed in *N. benthamiana* leaves. Extracted proteins were immunoprecipitated 36 hpi using anti-FLAG beads. The Co-IP proteins were detected by anti-T7 antibody. **(D)** Pth2 interacts with AvrPtoB and AvrPtoB^S335D^ *in vivo* by Co-IP assay. *35S:Pth2-T7*, *35S:AvrPtoB*-*FLAG*, *35S:AvrPtoB^S335D^-FLAG* or *35S:GFP-FLAG* were transiently expressed in *N. benthamiana* leaves. Extracted proteins were immunoprecipitated 36 hpi using anti-FLAG beads. The Co-IP proteins were detected by anti-T7 antibody. **(E)** AvrPtoB^S335D^ promotes Prf complex dissociation through Pth2 *in vivo* by Co-IP assay. *35S:Prf-Nterm*-*FLAG*, *35S:Prf-Nterm*-*HA, 35S:Prf-SCNL*-*HA, 35S:Pth2-T7*, *35S:GUS-T7*,and *35S:AvrPtoB* or *35S:AvrPtoBS^335D^* were transiently expressed in *N. benthamiana* leaves. Extracted proteins were immunoprecipitated 36 hpi using anti-FLAG beads. The Co-IP proteins were detected by anti-HA antibody. The band intensity was calculated by Image J. The smaller value represents the weaker interaction between proteins. **(F)** AvrPtoB^S335D^ induced cell death partially depends on Pth2. The transcription of *Pth2* was knocked down by VIGS in tomato. After 3 weeks, *Agrobacterium* GV2260 harboring *35S:AvrPtoB* and *35S:GFP* were infiltrated into tomato leaves. Ethanol was applied for decolonization after 72 hpi. The PCD area was quantified by image J. Value was means ± SD (n = 4 biological replicates; Student’s *t* test; \*\*\**p*< 0.001). All experiments were repeated three times with similar results.

### Pto-associated proteins originated from MLRs via the loss of the extracellular domain

To further trace the origin and evolution of Pto, we use a phylogenomic approach combining similarity search to identify proteins closely related to Pto from *S. pimpinellifolium,* across 141 plant species representing major plants diversity (Fig. 5; Supplementary Table S1). We found that Pto-associated proteins nest within the diversity of MLRs with strong support (ultrafast bootstrap approximation [UFBoot] = 99%). The lineage most closely related to Pto-associated proteins was HERK (HERCULES, one kind of CrRLK1Ls), which still retained a malectin-like domain (MLD). This result indicated that Pto-associated proteins originated from MLRs by losing their extracellular malectin domains (Supplementary Fig. S6; Supplementary Table S2). HERK occurred in Petunioideae, Solanoideae and Nicotianoideae, while Pto-associated proteins were present in Solanoideae and Nicotianoideae, but not in Petunioideae (Fig. 5; Supplementary Fig. S6). These findings showed that Pto-associated proteins that originated from an ancient duplication event occurred before the divergence of Petunioideae, Solanoideae and Nicotianoideae, approximately 60 million years ago during the Paleocene period, followed by the loss of the extracellular domain and gene loss in Petunioideae (Huang et al., 2023).

**Figure 5.**
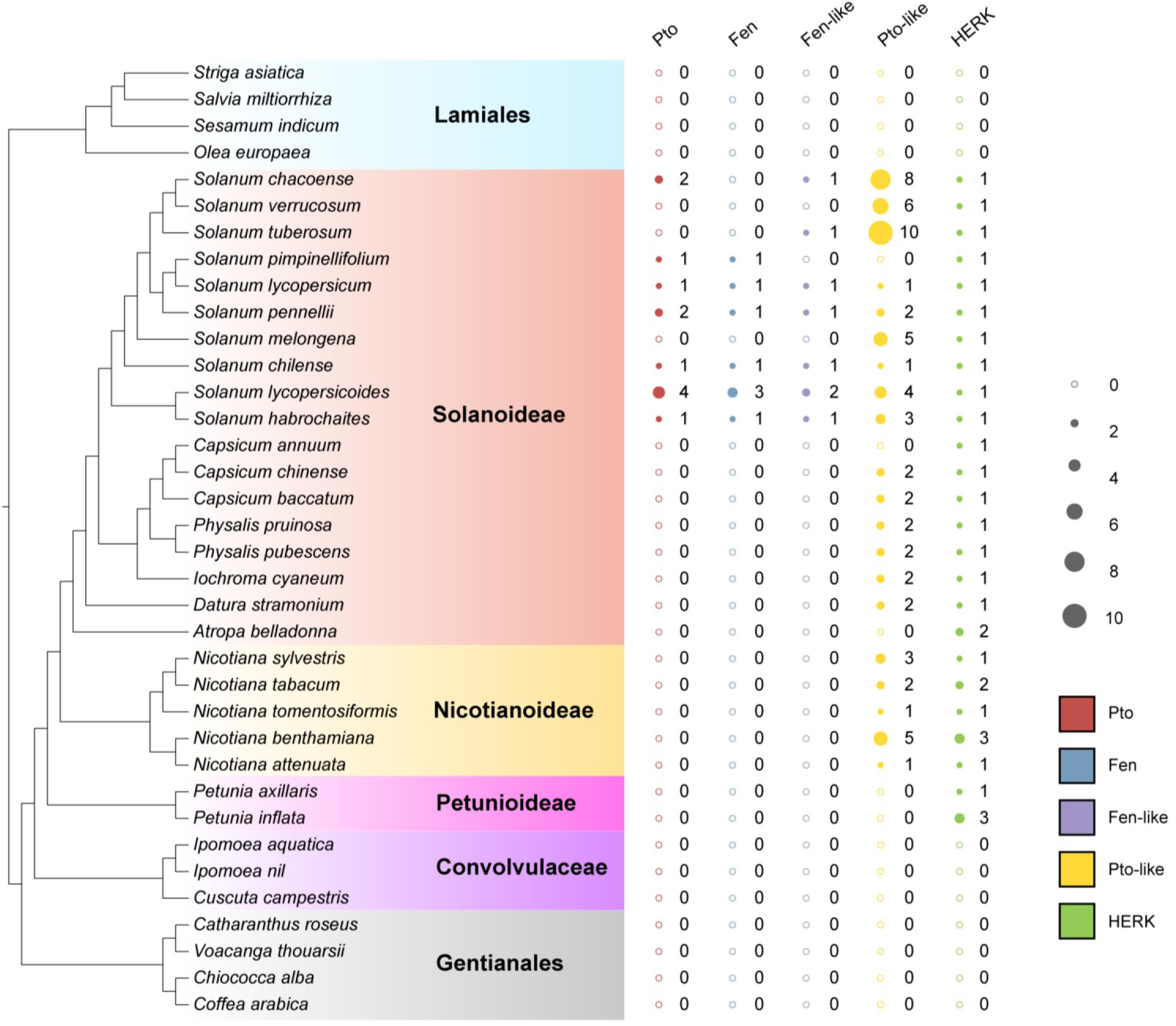
Distribution of Pto, Pto-like, Fen, Fen-like and HERK proteins in plants. For each plant species, the copy numbers of Pto, Pto-like, Fen, Fen-like and HERK were shown next to the species names. The solid and open circles represented the presence and absence of the corresponding proteins, respectively. The size of each solid circle reflected the copy number. The plant phylogeny was based on the literature (Leebens-Mack et al., 2019; Schwacke et al., 2025)

According to the phylogenetic relationship of Pto-associated proteins, Pto-associated proteins can be classified into at least nine major lineages (Pto-I-VII in *Solanum*, Pto-VIII-IX in Nicotiana), suggesting that these proteins have experienced at least six duplication events during the early evolution of *Solanum*, but were rarely duplicated in *Nicotiana* (Fig. 6). Fen and Pto homologs fell within the Pto-II and Pto-III lineages, respectively. Copy numbers of Fen and Pto proteins varied significantly among different species, indicating that these lineages have undergone frequent gene duplications and losses (Fig. 5; Fig. 6). Taken together, our results suggest that Pto-associated proteins originated from MLRs by losing their extracellular domains, at least before the split of Petunioideae, Solanoideae and Nicotianoideae, with Pto orthologs arising from recent duplications at the common ancestor of *Solanum*.

**Figure 6.**
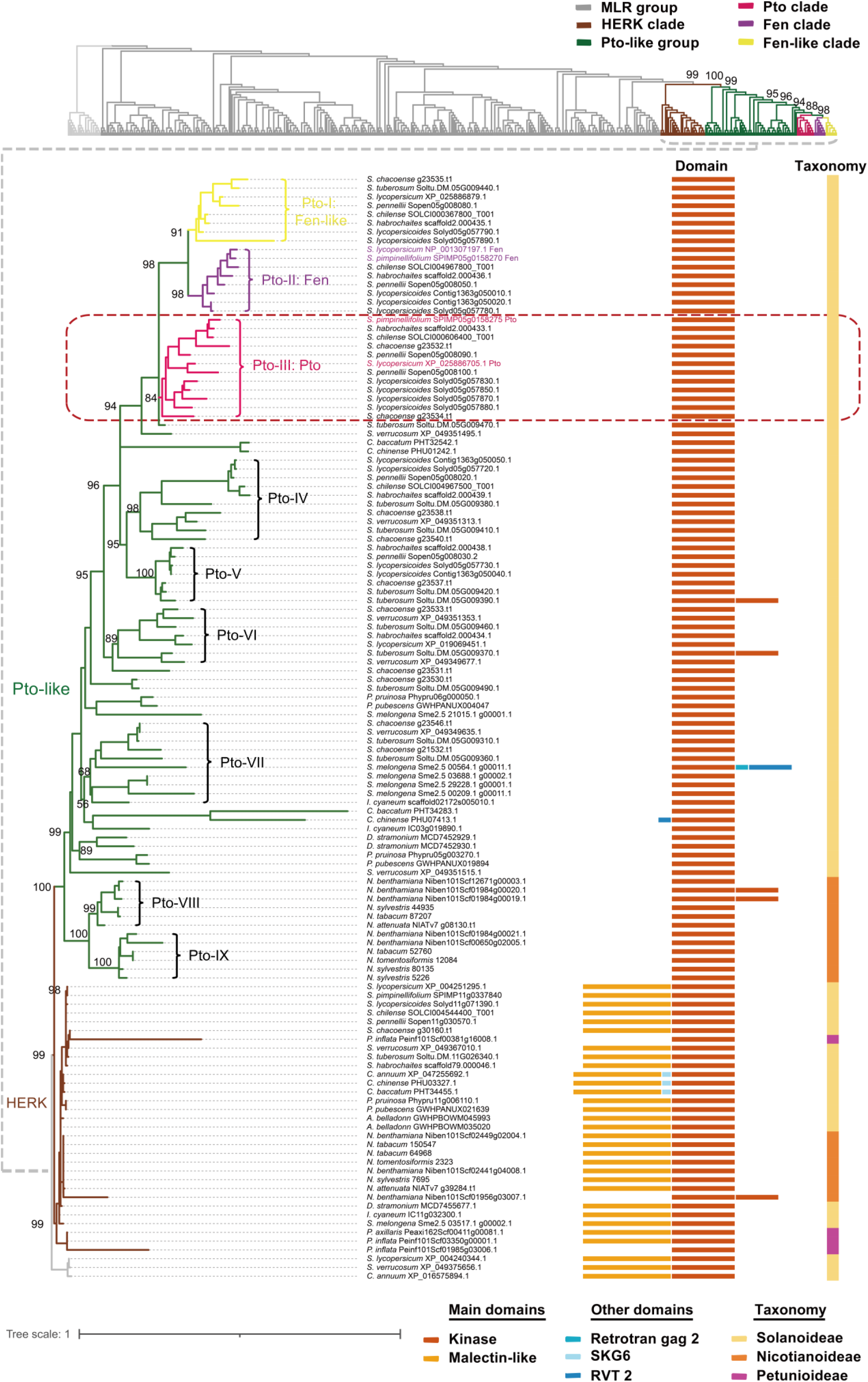
Phylogenetic relationship of Pto-associated proteins and HERKs. The phylogeny was reconstructed using the maximum likelihood method. Outgroups were indicated by light gray branches. The Pto-associated proteins were classified into four groups: Fen-like (in yellow branches), Fen (in purple branches), Pto (in pink branches), and Pto-like (in green branches). HERK proteins were indicated by gray branches. Pto group was highlighted by the red dashed line box. For each protein, the domain architecture was shown next to the protein name. Different domains were indicated using different color stripes, and the color key was shown on the bottom. The values near the nodes were UFBoot values. Retrotran_gag_2: gag-polypeptide of LTR copia-type retrotransposon. RVT 2: Reverse Transcriptase. SKG6: SKG6/AXL2 alpha-helix transmembrane domain. The detailed information of upper panel was shown in Supplemental Figure 6.

### SpHERK1s act as positive regulators of immunity against *Pst* DC3000

The phylogeny of Pto and MLRs from *S. lycopersicum* suggests that SpHERK1-1 (XP 004251295.1, HERK1), SpHERK1-2 (XP 004239762.1, HERK1) and SpHERK1-3 (XP 004240344.1, HERK1) are closely related to Pto (Supplementary Fig. S7A). Their intracellular domains share high similarity with Pto and Fen, including conserved ATP-binding sites and P+1 loops, which is an important surface for the interaction with AvrPto and AvrPtoB (Supplementary Fig. S7, B and C).

To validate the subcellular localization of SpHERK1-1, SpHERK1-2, and SpHERK1-3, we transiently expressed SpHERK1-1/-2/-3-GFP fusion proteins in *N. benthamiana* leaves via *Agrobacterium*-mediated transformation with plasma membrane marker AtPIP2-mCherry. Compared with GFP protein, SpHERK1s-GFP fusion protein co-localized with plasma membrane marker AtPIP2-mCherry at the plasma membrane (Supplementary Fig. S8A). Next, we analyzed the relative expression of three *SpHERK1* genes in RG-PtoR tomato after *Pst* DC3000 infection by RT-qPCR, and found that all *SpHERK1* genes were induced by *Pst* DC3000 treatment (Supplementary Fig. S8B). Then we used VIGS to knock-down three *SpHERK1s*, respectively (Supplementary Fig. S8C). Compared with the *TRV:EV* control plants, *SpHERK1s*-silenced tomatoes were more susceptible to *Pst* DC3000 infection (Supplementary Fig. S8D). Moreover, we generated two *SpHERK1-2* overexpressed tomato lines, OE-*SpHERK1-2-2*, and *OE-SpHERK1-2-14*. Compared with RG-PtoR tomato plants, the relative expression of *SpHERK1-2* was significantly upregulated in two transgenic lines (Supplementary Fig. S8E). Two *SpHERK1-2* overexpressed tomato lines were more resistant to *Pst* DC3000 infection (Fig. 7A). Taken together, these results indicated that three SpHERK1s are positive regulators in tomato disease resistance to *Pst* DC3000.

**Figure 7.**
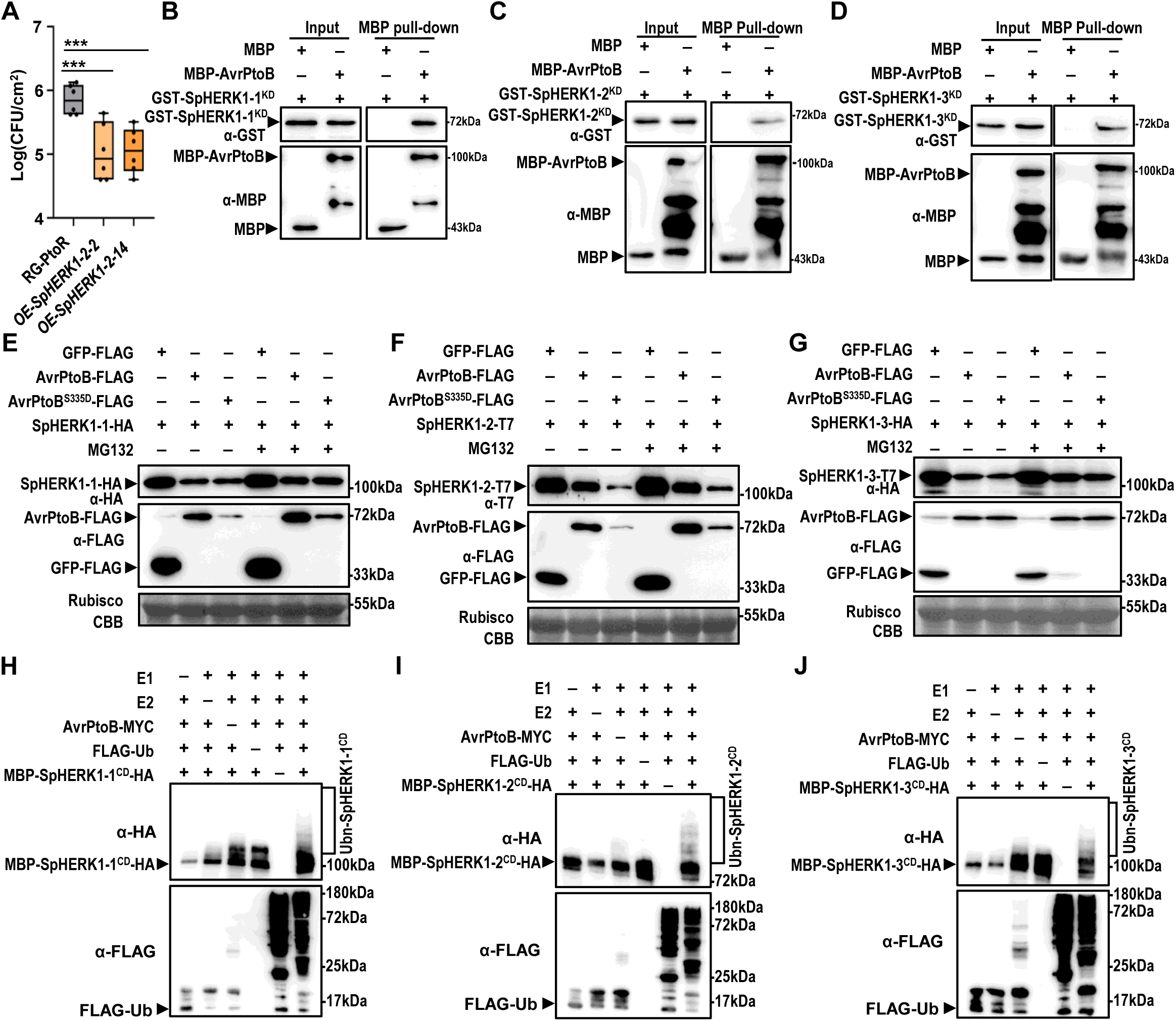
AvrPtoB interacts with SpHERK1s and targets SpHERK1s for degradation. **(A)** Growth curve analysis of *Pst* DC3000 in *SpHERK1-2* overexpressed tomato lines. Tomato plants were inoculated with *Pst* DC3000 at concentration of 2×10^8^ CFU/mL after 3-week-old. Plants were subjected to bacterial growth curve analysis at 3 dpi. Values were means ± SD (n = 6 biological replicates; Two-way ANOVA; \*\*\**p*< 0.001) **(B-D)** AvrPtoB interacts with SpHERK1-1 (**B**), SpHERK1-2 (**C**) and SpHERK1-3 (**D**) *in vitro* by MBP pull-down assay. The recombinant GST-SpHERK1-1^KD^, GST-SpHERK1-2^KD^ or GST-SpHERK1-3^KD^ and MBP-AvrPtoB were subjected to MBP pull-down assay. The interacting proteins were detected by immunoblot. **(E-G)** AvrPtoB and AvrPtoB^S335D^ degrade SpHERK1-1 (**E**), SpHERK1-2 (**F**) and SpHERK1-3 (**G**) via 26S proteasome in *N. benthamiana*. *35S:SpHERK1-1-HA*, *35S:SpHERK1-2-T7* or *35S:SpHERK1-3-HA* was transiently expressed in *N. benthamiana* leaves together with *35S:GFP-FLAG*, *35S:AvrPtoB-FLAG or AvrPtoB^S335D^-FLAG*. 100 μM MG132 was used to inhibit 26S proteasome-mediated protein degradation at 36 hpi. Samples were harvested at 6-8 h after MG132 treatment. The proteins were detected by immunoblot using corresponding antibody. CBB-stained Rubisco was used as loading control. **(H-J)** AvrPtoB ubiquitinates SpHERK1-1 (**H**), SpHERK1-2 (**I**) and SpHERK1-3 (**J**) *in vitro*. The bacterial lysates from *E. coli* BL21 strains expressing E1 (AtUBA1), E2 (AtUBC8), AvrPtoB-MYC, MBP-SpHERK1-2^CD^-HA, MBP-SpHERK1-2^CD^-HA or MBP-SpHERK1-3^CD^-HA and FLAG-Ub, or lacking one of these proteins were detected by immunoblot using anti-HA and anti-FLAG antibody.

### AvrPtoB degrades SpHERK1s through 26S proteasome pathway

Due to the similar spatial structures between Pto, Fen, and intracellular domain of SpHERK1s, we investigate whether AvrPtoB can interact with SpHERK1s. The MBP-pull-down, GST pull-down, and LCA assays showed that SpHERK1-1, SpHERK1-2, and SpHERK1-3 could interact with AvrPtoB *in vitro* and *in vivo* (Fig. 7, B to D; Supplementary Fig. S9, A to D). Co-expressing *AvrPtoB*, *AvrPtoB^S335D^*, *SpHERK1-1*, *SpHERK1-2*, and *SpHRERK1-3* in *N. benthamiana* leaves showed that AvrPtoB could degrade SpHERK1-1/2/-3 via 26S proteasome protein degradation pathway (Fig. 7, E to G). *In vitro* ubiquitination assays showed that AvrPtoB was able to ubiquitinate SpHERK1-1, SpHERK1-2, and SpHERK1-3 cytosolic domain (CD) (Fig. 7, H to J).

To determine whether SpHERK1-1/2/3 could phosphorylate AvrPtoB Ser335 site as Pto, we first used recombinant GST-SpHERK1-1^KD^, GST-SpHERK1-2^KD^, and GST-SpHERK1-3^KD^ to conduct the phosphorylation assay. Surprisingly, none of them could phosphorylate AvrPtoB. By Comparing the protein domain organization of Pto, SpHERK1-1, SpHERK1-2, and SpHERK1-3, we speculated that the N–terminal of Pto might also have important functions (Supplementary Fig. S9E). Then we cloned the full intracellular domain of SpHERK1-2 and performed phosphorylation assay. The results showed that MBP-SpHERK1-2^CD^ could phosphorylate AvrPtoB Ser335 site (Supplementary Fig. S9F). SpHERK1-1^CD^ and SpHERK1-3^CD^ can’t be purified through prokaryotic *E. coli* expression, so we expressed *35S:SpHERK1-1-HA* and *35S:SpHERK1-3-HA* in *N. benthamiana* leaves and purified them by anti-HA beads for phosphorylation assay. Consistent with SpHERK1-2, SpHERK1-1 and SpHERK1-3 could also phosphorylate AvrPtoB Ser335 site (Supplementary Fig. S9, G and H). These results indicated that SpHERK1-1/2/3 could interact with and phosphorylate AvrPtoB as Pto.

### Arg158 and Glu258 residues of Pto protect it from being degraded by AvrPtoB

To further elucidate the molecular mechanism by which AvrPtoB can degrade Pto-related MLRs SpHERK1-1, SpHERK1-2, and SpHERK1-3, but not Pto. We conducted a conservation analysis of protein sequences. Conserved sequences with 50% conservation in the Pto-associated proteins and HERK-clade proteins were identified respectively. The alignment result showed that Pto 158 and 258 sites were conserved (50%) lysine sites in the HERK-clade proteins, but lost high conservation (50%) in Pto-associated proteins (Supplementary Fig. S10A). Because canonical ubiquitination occurs on lysine residues, we inferred that these sites might have biochemical significance. We mutated Arg158 and Glu258 of Pto to lysine (R158K and E258K) and examined the effects of these two mutations on Pto stabilization. We transiently co-expressed *35S:Pto-HA*, *35S:Pto^R158K^-HA*, *35S:Pto^E258K^-HA* and *35S:Pto^R158KE258K^-HA* with *35S:GFP-FLAG* or *35S:AvrPtoB-FLAG* in *N. benthamiana* leaves. As shown in Supplementary Fig. S10B, compared with Pto, the protein bands of Pto^R158K^, Pto^E258K^, PtoR^158KE258K^ were significantly reduced in the presence of AvrPtoB. To investigate whether these mutations affect the kinase activity of Pto, we constructed GST-Pto^R158K^, GST-Pto^E258K^ and performed kinase assay in vitro. Phosphorylation assay result demonstrated that mutations at these two sites didn’t affect the phosphorylation ability of Pto at AvrPtoB Ser335 (Supplementary Fig. S10C). These data suggest that residues Arg158 and Glu258 of Pto are crucial for avoiding Pto from being degraded by AvrPtoB.

## Discussion

In this study, we revealed a novel mechanism by which phosphorylation of a bacterial effector to initiate a stronger immunity, and elucidated the key evolutionary adaptations of the first identified disease-resistance protein Pto (Fig. 8).

**Figure 8.**
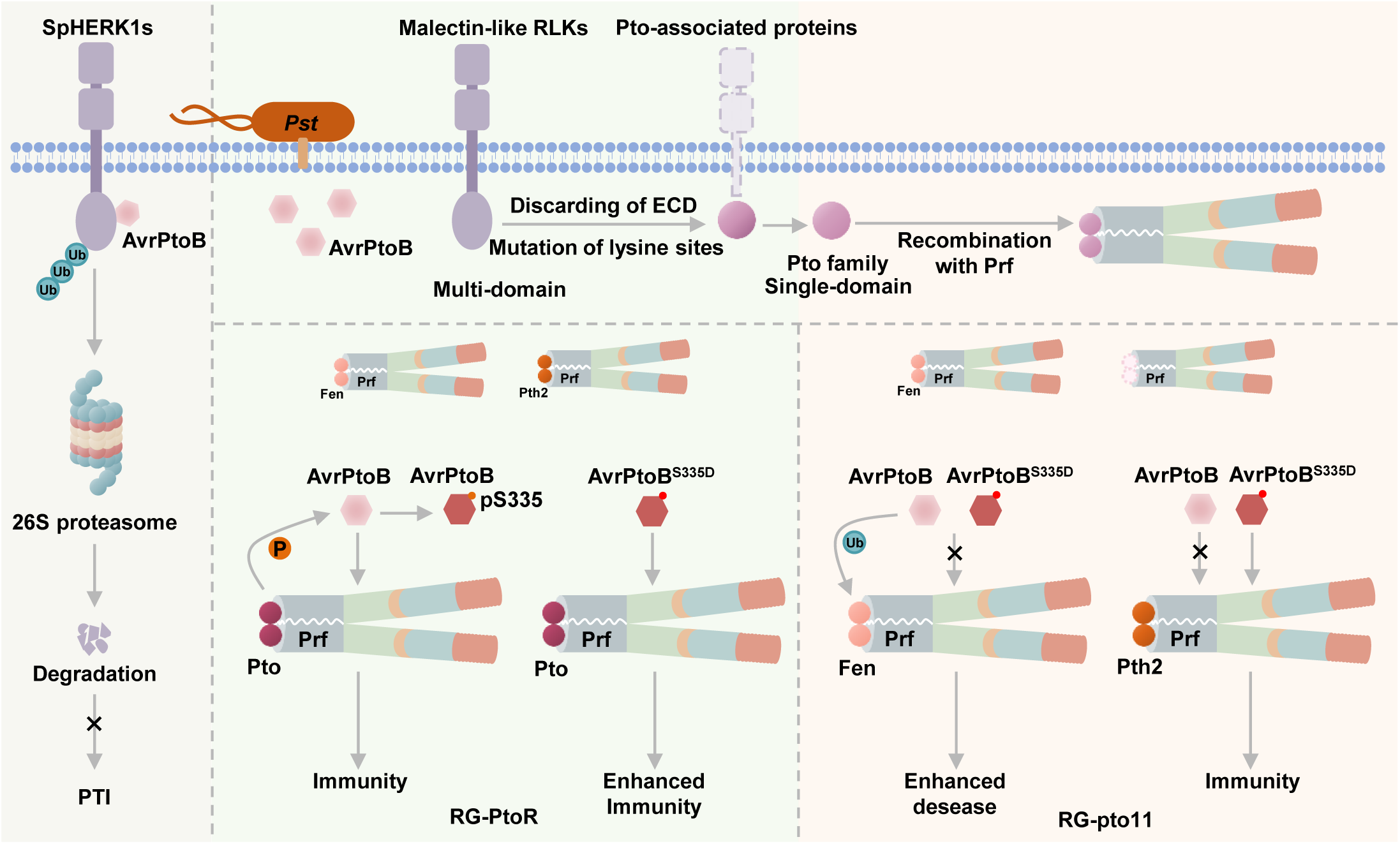
Working model of the Pto evolution and Pto, HERK1s–mediated immune response. *Pst* secretes effector–AvrPtoB into host plant, which can ubiquitinate and degrade SpHERK1s through 26S proteasome pathway, thereby inhibits plant PTI response. Under long-term selection pressure, plant malectin-like RLKs evolved into Pto-associated proteins by losing their extracellular domain. Among them, Pto and its homologous proteins including Fen and Pth2 can interact with Prf Nterm domain to form resistance complex. In wild-type tomato plant (RG-PtoR), Pto phosphorylates AvrPtoB at Ser335 site, which enhances the Prf-mediated immune response. In pto mutant (RG-pto11), AvrPtoB (orange dot) and AvrPtoB^S335D^ (mimic phosphorylation of AvrPtoB Ser335 site, red dot) ubiquitinate Fen, so they can’t activate Prf-mediated immune response through Fen. While AvrPtoB^S335D^ but not AvrPtoB can activated Prf-mediated immune response through Pth2.

In the interaction between plants and pathogens, pathogenic bacteria secrete effector proteins into the host plant through the type III secretion system to suppress plant immunity and maintain their colonization. AvrPtoB is a well-studied E3 ubiquitin ligase (Rosebrock et al., 2007). In *Arabidopsis*, no NLR protein can recognize AvrPtoB and trigger ETI, consequently, AvrPtoB inhibits plant PTI by degrading multiple disease-related proteins, such as FLS2, CERK1, and LecRK-IX.2 (Rosebrock et al., 2007; Göhre et al., 2008). Plants have also evolved mechanisms to suppress the function of AvrPtoB, for example, LecRK-IX.2 phosphorylates AvrPtoB at Ser335 site, which can disrupt its self-association of AvrPtoB and weaken its virulence (Xu et al., 2020). However, in tomato, AvrPtoB acts as an avirulence factor and is recognized by a resistant complex composed of Pto, Prf and NRCs, thereby elevating the immune response to the stronger ETI level.

Upon recognition of AvrPtoB, the Pto/Prf complex dissociates, activating downstream immune responses (Sheikh et al., 2023). Pto, known as a cytoplasmic receptor kinase, phosphorylates the Thr450 site of AvrPtoB. Phosphorylation at Thr450 attenuates AvrPtoB’s activity and reduces its pathogenicity in RG-pto11 tomato plants (Ntoukakis et al., 2009). Our study found that Pto can also phosphorylate the Ser335 site of AvrPtoB (Fig. 1A). PCD assays revealed that the phosphomimetic AvrPtoB^S335D^ induces PCD in RG-PtoR and RG-pto11 tomato plants but not in RG-prf3, indicating that the PCD triggered by AvrPtoB^S335D^ is dependent on Prf but not entirely on Pto (Fig. 2A). Growth curve results showed that both bacterial strains maintained similar growth levels in RG-PtoR, suggesting that AvrPtoB may exist in a fully phosphorylated state at the Ser335 site in tomatoes, making phosphorylation at this site critical for downstream immune recognition (Fig. 2, B and C). However, compared to AvrPtoB^WT^, the strain expressing AvrPtoB^S335D^ exhibited significantly reduced pathogenicity in RG-pto11 tomato plants, implying that in *pto* mutants, other proteins may cooperate with Prf to recognize AvrPtoB^S335D^.

NLR proteins play a pivotal role in plant immunity by recognizing pathogen effectors. In most cases, after recognition of pathogen effectors, NLR forms higher-order protein complexes known as resistosomes to integrate signaling through Ca^2+^ permeable channels and trigger downstream defense mechanisms (Wang et al., 2019; Bi et al., 2021; Förderer et al., 2022; Liu et al., 2024a; Madhuprakash et al., 2024). However, some NLR proteins exist as complexes in the absence of effector proteins, such as NRC2 (SlNRC2) in *S. lycopersicum* and NbNRC2 in *N. benthamiana.* The dimerization and oligomerization of SlNRC2 not only stabilize the inactive state but also sequester SlNRC2 from assembling into an active form (Ma et al., 2024). NbNRC2 accumulates *in vivo* as a homodimer and forms a higher-order oligomer upon activation by its upstream virus disease resistance protein Rx (Selvaraj et al., 2024). For the Prf/Pto complex, the N-terminal domain of Prf facilitates the oligomerization of itself and mediates its interaction with Pto, thereby positioning two Pto molecules in close proximity while maintaining them in an inactive conformational state in the absence of effector proteins (Sheikh et al., 2023). Following AvrPto/AvrPtoB effector recognition, the Prf/Pto complex dissociates into distinct modules. Our results showed that phosphorylation of AvrPtoB at Ser335 promotes dissociation of Prf/Pto resistance complex and is a conserved function among diverse Pto variants (Fig. 1B; Supplementary Fig. 3). Previous study has shown that Pto acts as a decoy to recognize AvrPtoB via its P+1 loop structure. Disrupting this structure can activate the Prf-mediated immune response (Ntoukakis et al., 2013). We speculated that AvrPtoB ^S335D^ had a stronger ability to disrupt this structure of Pto compared to AvrPtoB. This conformational change induces trans-phosphorylation within Pto, which subsequently serves as an activating signal and triggers the dissociation of the complex.

Although AvrPtoB-triggered dissociation of the Prf/Pto complex depends on Pto, AvrPtoB^S335D^ could still induce PCD in RG-pto11 lines. Our further research demonstrates that Pth2, one of Pto homologs, forms complex with Prf and mediates AvrPtoB^S335D^-induced PCD in RG-pto11 lines (Fig. 4). The residual cell death in Pth2 VIGS lines was due to incomplete silencing of Pth2 in RG-PtoR, or other unidentified components. These results indicate that tomato has evolved multiple Pto homologs to guarantee the immunity activation of Prf complex after recognition of AvrPtoB.

Previous studies have suggested that plant RLKs might have evolved in a modular fashion through frequent domain gains or losses, and that novel immunity modules might have originated from preexisting proteins via molecular tinkering (Gong and Han, 2021; Gong et al., 2022). In plants, more than 10 distinct extracellular domain (ECD) structures are associated with the kinase domain, such as LRR-, LysM-, Malectin-, G-lectin-, L-lectin-, Duf26-, and WAK-ectodomains. In *Arabidopsis*, to initiate defense, NLR protein HOPZ-ACTIVATED RESISTANCE 1 (ZAR1) and RLCK protein HOPZ-

ETI-DEFICIENT 1 (ZED1)-related kinases (ZRKs) can form a resistosome to perceive effector-induced perturbations indirectly (Wang et al., 2019). Largescale phylogenetic analysis shows that ZRKs originated from wall-associated protein kinases (WAKs) via the loss of the extracellular domain before the split of eudicots and monocots during the Jurassic period (Gong et al., 2022). Interestingly, we found that Pto-associated proteins originated from MLRs (CrRLK1L) via the loss of the extracellular domain, supporting the molecular tinkering model. Discarding of ECD simplified the kinase interacting with AvrPtoB-like proteins during the Paleocene period. It allowed the simplified kinase (Pto) to leave the cell membrane and gain the capacity to signal via Prf, which perhaps would not have been possible in the context of a full-length RLK. In addition, Pto-associated proteins were distributed only in Solanales (Fig. 5; Supplementary Fig. S6), indicating the potential similar immunity functions of Pto-associated proteins from other Solanales species, which remains to be further validated.

Lysine residues are key sites for 26S proteasome pathway. In *Arabidopsis*, there are three similar ADR1 class helper NLRs (ADR1, ADR1-L1, and ADR1-L2). Wang et al. show that AvrPtoB degrades ADR1-L1 and ADR1-L2, however, ADR1 evades degradation by diversifying the ubiquitination sites of AvrPtoB. Mutation of two key sites in ADR1, with E35K/R49K substitutions, promoted the ubiquitination of ADR1 by AvrPtoB (Wang et al., 2023). Recent studies suggest that AvrPtoB interacts with transcription factor ELONGATED HYPOCOTYL 5 (HY5) and promotes its ubiquitination and degradation. HY5 variants carrying lysine-to-arginine substitutions exhibit significantly reduced ubiquitination (Liu et al., 2025). These results indicate that lysine residues are key sites for AvrPtoB-medicated 26S proteasome pathway. Although Pto, SpHERK1-1, SpHERK1-2, and SpHERK1-3 all could phosphorylate AvrPtoB at Ser335, AvrPtoB degrades SpHERK1s through 26S proteasome pathway, in contrast, Pto is more resistant to the degradation of AvrPtoB (Fig. 1A; Supplementary Fig. S9, F to H; Fig. 7, E to G). Our conservation analysis of protein sequences identified two important residues (Arg158 and Glu258) of Pto, which were conserved lysines in HERK proteins, and protected Pto from degradation (Supplementary Fig. S10A). If these two sites are mutated to lysine, the degree of Pto degradation by AvrPtoB is greatly enhanced (Supplementary Fig. S10B). This result indicates that Solanales plants have been continuously adapting to the stress of AvrPtoB-like proteins since at least the Paleocene. Therefore, for further molecular breeding and crop engineering, modifying the key lysine of some resistant proteins is a superior, naturally selected strategy.

## Materials and methods

### Plant materials and growth conditions

Tomato Rio Grande (RG)-PtoR, RG-pto11, RG-prf3, 1 cultivated tomato (*Solanum lycopersicum* MoneyMaker) and 9 wild tomatoes (*S. lycopersicoides*, *S. habrochaites*, *S. pennellii*, *S. chilense*, *S. peruvianum*, *S. corneliomulleri*, *S. neorickii*, *S. chmielewskii*, *S. galapagense*) were obtained from Tomato Genetics Resource Center (TGRC, https://tgrc.ucdavis.edu/). The *SpHERK1-2* coding sequence was amplified and cloned into the pCAMBIA1305 vector. The recombinant vector was transformed into *Agrobacterium tumefaciens*. The resulting strain was used to transform the RG-PtoR tomato plants. Tomato and *Nicotiana benthamiana* plants were grown at 25°C under 10 h light/14 h dark conditions for 3-4 weeks.

### *In Vitro* Kinase assay

The *in vitro* kinase assays were performed according to the method described by Xu et al. (Xu et al., 2020). Protein expression and purification were performed according to the method described by Luo et al. (Luo et al., 2017). For *in vitro* kinase assay, purified proteins including 2 μg kinases and 2 μg substrates were added into 30 μL kinase buffer containing 20 mM Tris-HCl (pH7.5), 10 mM MgCl_2_, 1 mM CaCl_2_, 1 mM DTT, 1 mM cold ATP and incubated at 30°C for 45 min. Protein samples were boiled for 10 min after mixing with 3× laemmli buffer (137.5 mM Tris-HCl (pH 6.8), 22.5% Glycerol, 6.6% SDS, 0.02% BPB, 300 mM DTT) and separated by 8% SDS-PAGE. Phosphorylated proteins were analyzed by immunoblot using anti-pSer/Thr antibody (ECM Bioscience, PP2551, 1:10000) and anti-pS335 antibody (1:200) prepared by Abmart (Xu et al., 2020). Inputs of all proteins were detected by anti-MBP (EASYBIO, BE2021-100, 1:10000), anti-GST (Transgen Biotech, HT601-01, 1:10000) and anti-HIS (Sigma-Aldrich, SAB1305538, 1:10000) antibodies. For ^32^P isotope-labeled reactions, 30 μL kinase reaction mixture contained 25mM Tris-HCl (pH7.5), 10 mM MgCl_2_, 10 mM MnCl_2_, 1 mM DTT, 50 μM cold ATP, 1 μCi ^32^P-ATP, 2 μg kinases and 2 μg substrates and incubated at 30°C for 45 min. Denatured proteins were separated on 10% SDS-PAGE, and phosphorylated proteins were visualized by X-ray film exposure.

For the SpHERK1-1-HA and SpHERK1-3-HA kinase assay, *35S: SpHERK1-1-HA* and *35S: SpHERK1-3-HA* were expressed in *Nicotiana benthamiana* leaves and enriched using anti-HA agarose beads, respectively. Then the SpHERK1-1-HA and SpHERK1-3-HA were respectively added to kinase buffer containing 2 μg of substrate and incubated at 30°C for 45 min. Protein samples were boiled for 10 min after mixing with 3× laemmli buffer, and separated by 8% SDS-PAGE. Phosphorylated proteins were analyzed by immunoblotting with anti-pS335 antibody. Primers used are listed in Supplementary Table S3.

### Phylogenetic analyses

To investigate the distribution and evolution of Pto across plants, similarity searches and phylogenetic analyses were conducted on 141 representative plant species (Supplementary Table S1). First, the BLASTp algorithm was used to search plant proteomes with the serine/threonine kinase catalytic (STKc) domain sequence of Pto (*S. pimpinellifolium* Pto) as the query, with an e-value cutoff of 10^−5^. All the kinase domain sequences of significant hits were aligned with Pto STKc domain using MAFFT (Katoh and Standley, 2013; Gong and Han, 2021). The alignments were refined using the R pipline (sites with less than 98% blank were retained). Large-scale phylogenetic analyses were performed using the approximate maximum-likelihood method implemented in FastTree (Price et al., 2010). Significant hits that clustered with Pto were retrieved as Pto-related proteins (Supplementary Table S2). The domain organization of each protein was annotated by CD-Search (Lu et al., 2020).

To explore the evolutionary relationships among Pto-related proteins, a phylogenetic tree was reconstructed. The sequences were aligned with the Pto kinase domain of Pto-related proteins and representative eukaryotic protein kinases (ePKs) were aligned using the L-INS-I strategy in MAFFT (Katoh and Standley, 2013). The representative eukaryotic proteins were selected based on the aforementioned large-scale phylogenetic tree. The alignments were refined using the R pipline (sites with less than 90% blank were retained). Phylogenetic analyses were performed using the maximum likelihood method implemented in IQ-TREE (Nguyen et al., 2015). The ModelFinder algorithm was used to choose the best-fit substitution model (Kalyaanamoorthy et al., 2017). Supports for the nodes were assessed using the UFBoot approach with 1,000 replicates (Hoang et al., 2018). Related phylogenetic trees were annotated using iTOL (Letunic and Bork, 2019). Pto-related proteins were then counted across 141 plant species. Species without Pto-related proteins were not shown (Fig. 5). To compare the conservation patterns between proteins in the HERK clade and Pto-associated clade, sequences of the HERK and Pto-associated clades were extracted from aforementioned aligned sequences of Pto-related proteins. Extracted sequences were refined manually (only Pto sites were retained). A R pipline was used to identify conservative sequences of HERK and Pto-associated clades, respectively (50% conservation). Pto kinase sequence, conserved sequences of HERK, and Pto-associated clades were compared with GeneDoc program (Nicholas, 1997).

To identify the homologous RLCKs and MLRs closely related to Pto in *S. lycopersicum*, all the aforementioned significant hits from *S. lycopersicum* proteomes were aligned with Pto STKc domain using MAFFT (Katoh and Standley, 2013; Gong and Han, 2021). Large-scale phylogenetic analyses were also performed using FastTree (Price et al., 2010). The domain organization of each protein was annotated using CD-Search (Lu et al., 2020), and significant RLCKs and MLRs hits that clustered with Pto were retrieved as Pto-clustered proteins. To further explore the phylogenetic relationships among Pto-clustered proteins, a phylogenetic tree was reconstructed. The sequences aligned with Pto kinase domain of the Pto-clustered proteins and representative protein kinases were aligned using MAFFT (Katoh and Standley, 2013). The representative proteins were selected based on the aforementioned large-scale phylogenetic tree. Phylogenetic analyses were then performed using IQ-TREE as described above (Nguyen et al., 2015; Kalyaanamoorthy et al., 2017; Hoang et al., 2018). The resulting phylogenetic tree was annotated using iTOL (Letunic and Bork, 2019).

### Tomato leaves cell death assay

*Agrobacterium*-mediated transient expression in tomato leaves was performed as described previously (Janjusevic et al., 2006). Recombinant vectors *35S:AvrPtoB-T7*, *35S:AvrPtoB^S335D^-T7*, *35S:AvrPtoB^S335A^-T7* and *35S:AvrPtoB^F479A^-T7* were transformed into *A. tumefaciens* GV2260, which was syringe-infiltrated into tomato leaves at a final OD_600_ of 0.075. After 3 days of inoculation, the leaves were decolorized with anhydrous ethanol and the PCD was detected at the injection site. Primers used are listed in Supplementary Table S3.

### Bacterial strains construction

The *Pst* ΔavrPtoΔavrPtoB mutant strain was constructed according the method described by Wei (Wei et al., 2013). Two 500 bp left and right-flanking regions of the target gene were overlapped and cloned into pKMS1. The recombinant vector was transformed into *Pst* by triparental mating. The deletion mutant was obtained by sucrose counter-selection and PCR verification. To construct the AvrPtoB and its point mutant complementary strains, the *AvrPtoB* and its point mutant were cloned into pBBR-MCS2. The recombinant vectors were transformed into *Pst* ΔavrPtoΔavrPtoB mutant strain by electroporation (Xu et al., 2020).

### Pathogen infection assay

*Pst* DC3000 was grown on NYGA medium (5 g/L Peptone, 3 g/L Yeast extract, 2% Glycerol and 15 g/L Agar) at 28°C for 2 days. For syringe infection, the pathogen was suspended in a 10 mM MgCl_2_ solution and adjusted 5×10^4^ CFU/mL. Four-week-old tomato leaves were injected with the pathogen, and bacterial growth curve was calculated after 3 days infection. For spray inoculation, the pathogen was suspended in a 5 mM MgCl_2_ solution containing 0.025% Silwet L-77 and adjusted 2×10^8^ CFU/mL. Four-week-old tomato plants were soaked in pathogen solution for 1 min and the growth curve was calculated after 5 days infection.

### MBP pull-down assay

3 μg MBP-tagged protein and 3 μg GST-tagged protein were added to 1mL TEN buffer (20 mM Tris-HCl (pH7.5), 0.1 mM EDTA (pH8.0), 100 mM NaCl and 0.2% Triton X-100) containing 20 μL Amylose Resin (NEB, E8021V) or Glutathione Sepharose 4B (Cytiva, 17075601) and incubated at 4°C for 1.5 h under gentle rotation. Then beads were washed 5-7 times with 1 mL NTEN buffer (20 mM Tris-HCl (pH 7.5), 0.1mM EDTA (pH 8.0), 300 mM NaCl, 0.5% NP-40). 80 μL 1× laemmli buffer were added into beads and boiled for 10 min to release the bound proteins. Proteins were separated by SDS-PAGE and detected by immunoblot using anti-GST (Transgen Biotech, HT601-01, 1:10000) and anti-MBP (EASYBIO, BE2021-100, 1:10000) antibodies.

### Split-luciferase complementation assays in *N. benthamiana* leaves

The coding DNA sequences of detected genes were cloned into pCAMBIA1300-nLUC and pCAMBIA1300-cLUC, respectively. Recombinant plasmids were then transformed into *Agrobacterium* EHA105. The bacterial solution was injected into 4-week-old *N. benthamiana* leaves. The inoculated leaves were infiltrated with 0.5 mM luciferin at 36 hpi and kept in the dark for 5 min. Luciferase images were captured using the cooled CCD imaging apparatus. Primers used are listed in Supplementary Table S3.

### *In Vitro* ubiquitination assay

For the *in vitro* ubiquitination assay in *E. coli*, *SpHERK1-1^CD^*, *SpHERK1-2^CD^*, *SpHERK1-3^CD^*, *Pth2*, *Fen*, *Pto* and *AvrPtoB* genes were cloned into pCDFDuet or pACYCDuet vectors, respectively. Different combinations of plasmids that could express different transform ubiquitination components were transformed into *E. coli* BL21. When the bacteria solution reached OD_600_=0.4-0.6 at 37°C, the expression of target protein was induced by addition of 0.5 mM IPTG. The bacteria were cultured at 25°C for 10-12 h and kept at 4°C overnight. Bacteria were collected from 500 μL solution by centrifugation, resuspended with 100 μL 1× laemmli buffer and boiled for 10 min. The crude extracts were separated by SDS-PAGE and analyzed by immunoblotting with the corresponding antibodies.

For the *in vitro* ubiquitination assay by purified recombinant proteins, 30 μL reaction mixture contained 50 mM Tris-HCl (pH 7.5), 2 mM ATP, 5 mM MgCl_2_, 2 mM DTT, 1.25 μg FLAG-tagged ubiquitin, 250 ng His-E1 (wheat), 500 ng His-E2 (Human UbcH5b), 100 ng or 500 ng MBP-AvrPtoB or its mutants, and 500 ng GST-Fen, and incubated at 30°C, 900 rpm for 2 h. Protein samples were boiled for 10 min after mixing with 3× laemmli buffer, and separated by 8% SDS-PAGE. The proteins were analyzed by immunoblotting using indicated antibodies. Primers used are listed in Supplementary Table S3.

### Virus-induced gene silencing in tomato plants

Specific sequences of *SpHERK1-1*, *SpHERK1-2*, and *SpHERK1-3* genes were cloned into the pTRV2 vector and transformed into *Agrobacterium* EHA105. Bacteria containing recombinant pTRV2 and pTRV1 vector, respectively, were resuspended with infiltration buffer (10 mM MgCl_2_, 10 mM MES (pH5.6) and 100 μM acetosyringone) and mixed in equal volume. Bacterial solution was infiltrated into 2-week-old tomato leaves. The bacteria containing the *PDS* gene were used as control. When control tomato leaves showed bleaching phenotype, the corresponding genes expression levels in all plants were detected, and the plants with high silencing efficiency were selected for *Pst* DC3000 resistance detection. Primers used are listed in Supplementary Table S3.

### Total RNA extraction and RT-qPCR analysis

Total RNA was isolated from 4-week-old tomato plants using TRIzol reagent (TIANGEN Biotech Co., LTD) according to the method provided in the instructions. 1 μg total RNA was reverse-transcribed to synthesize first-strand cDNA using the HiScript II Q RT SuperMix for qPCR (+gDNA wiper) kit. RT-qPCR was performed using Taq Pro Universal SYBR qPCR Master Mix on a QuantStudio 1 Real-Time PCR System. Tomato *Actin* was used as reference gene. Primers used are listed in Supplementary Table S3.

### Subcellular localization in *N. benthamiana* leaves

*SpHERK1-1*, *SpHERK1-2*, and *SpHERK1-3* genes were cloned into a pMD1-GFP vector, which was then transformed into *Agrobacterium* EHA105. The Bacterial solution was infiltrated into 3-week-old *N. benthamiana* leaves. Images were obtained using a Lecia SP8 confocal laser microscope after 48 hpi. The *35S:PIP2A-mCherry* was used as cell membrane marker. The excitation wavelengths of GFP and mCherry were 488 nm and 552 nm, respectively. Primers used are listed in Supplementary Table S3.

### Co-immunoprecipitation assay

*Agrobacterium* EHA105 strains containing the indicated recombinant plasmids were infiltrated into 4-week-old *N. benthamiana* leaves. 1 g of inoculated leaves were harvest at 2 dpi, and total protein was extracted using 2 mL extraction buffer (50 mM Tris-HCl (pH 7.5), 150 mM NaCl, 0.1% Triton X-100, 0.2% NP-40, 1×protease inhibitor). Then the total protein was incubated with 20 μL anti-FLAG M2 Affinity Gel (Sigma-Aldrich, A2220) at 4°C for 1.5 h under gentle rotation. After that, the beads were washed 6 times using extraction buffer and boiled for 10 min to release proteins. The indicated proteins were separated by SDS-PAGE and analyzed by immunoblotting with the anti-FLAG (Sigma-Aldrich, F3165, 1:10000), anti-HA (Cell Signaling Technology, 3724S, 1:10000) and anti-T7 (Abcam, ab9115, 1:10000) antibodies. Primers used are listed in Supplementary Table S3.

### Statistical analysis

All statistical analysis of most data was performed using GraphPad Prism 9.0.0. Significant difference was calculated using two-way ANOVA or two-tailed Student’s *t* test.

## Accession numbers

Sequence data from this article can be found in the NCBI data libraries under accession numbers: Pto (AAF76306), Fen (AAF76307), SlPto (NP_001307158), Prf (AAF76308), AvrPtoB (WP_011104378), Pth2 (AAF76305), Pth3 (AAF76304), SpHERK1-1 (XP_004251295.1), SpHERK1-2 (XP_004239762.1), SpHERK1-3 (XP_004240344.1). The Pto homologs sequences can be found in published tomato genome sequences (http://caastomato.biocloud.net) under accession numbers: SlycPto (Slyd05g006930.1), ShabPto (Shab05g007000.1), SpenPto (Spen05g008090.1), SchiPto (Schi05g007810.1), SperPto (Sper05g007200.1), ScorPto (Scor05g007100.1), SneoPto (Sneo05g006760.1), SchmPto (Schm05g006950.1), SgalPto (Sgal05g007630.1).

## Acknowledgments

The authors thank Prof. Qian Chen (China Agricultural University, China) for technical assistance with the E3 ligase activity assay, Dr. Na Jiang and Dr. Linlu Qi (China Agricultural University, China) for technical support. The authors also thank the High-performance Computing Platform of the China Agricultural University. No conflict of interest is declared.

## Author contributions

N.X., J.L., L.L., X.Z., and Z.G. conceived, conceptualized, and designed the experiments. L.L., X.Z., and Z.G. performed most of the experiments. J.S., Q.C., W.Wu., J.Y., and W.Wang. helped with the molecular cloning, phenotype statistics, and data analysis. N.X., L.L, X.Z., and Z.G. wrote the manuscript.

## Supplementary data

The following materials are available in the online version of this article.

**Supplementary Figure S1.** Kinase-dead mutants of Pto, SlPto and Fen can’t phosphorylate AvrPtoB at Ser335.

**Supplementary Figure S2.** Multiple sequence alignment of Pto and its homologous proteins from wild and cultivated tomato species.

**Supplementary Figure S3.** Pto homologs phosphorylate AvrPtoB at Ser335.

**Supplementary Figure S4.** AvrPtoBS335D doesn’t dissociate Prf complex through Fen.

**Supplementary Figure S5.** AvrPtoBS335D induces the dissociation of Prf complex through Pth2 but not Pth3.

**Supplementary Figure S6.** Phylogenetic analysis of the tomato Pto-related proteins.

**Supplementary Figure S7.** Homology analysis of Pto and tomato MLR sequence.

**Supplementary Figure S8.** Three SpHERK1 proteins positively regulate resistance to Pst DC3000 in tomato.

**Supplementary Figure S9.** SpHERK1s interact with and phosphorylate AvrPtoB Ser335 site.

**Supplementary Figure S10.** The amino acid sites Arg158 and Glu258 are important for the stability of Pto.

**Supplementary Table S1.** Sources of the proteomes.

**Supplementary Table S2.** Domain architectures.

**Supplementary Table S3.** Primers used in this study.

## Funding

This study was supported by the Beijing Natural Science Foundation (6262014), the National Key Research and Development Program of China (2024YFD1200602), the National Natural Science Foundation of China (grants No. 32270297 and 32225043), and the 2115 Talent Development Program of China Agricultural University.

## Data availability

All data supporting the conclusions of the article are available in the main text and Supplementary Information.

## Notes

### Competing Interest Statement

The authors have declared no competing interest.

## References

Abramovitch, R.B., and Martin, G.B. (2005). AvrPtoB: a bacterial type III effector that both elicits and suppresses programmed cell death associated with plant immunity. FEMS Microbiol Lett 245, 1–8.

Abramovitch, R.B., Janjusevic, R., Stebbins, C.E., and Martin, G.B. (2006). Type III effector AvrPtoB requires intrinsic E3 ubiquitin ligase activity to suppress plant cell death and immunity. Proc Natl Acad Sci U S A 103, 2851–2856.

Abramovitch, R.B., Kim, Y.J., Chen, S., Dickman, M.B., and Martin, G.B. (2003). Pseudomonas type III effector AvrPtoB induces plant disease susceptibility by inhibition of host programmed cell death. Embo j 22, 60–69.

Bi, G., Su, M., Li, N., Liang, Y., Dang, S., Xu, J., Hu, M., Wang, J., Zou, M., Deng, Y., Li, Q., Huang, S., Li, J., Chai, J., He, K., Chen, Y.H., and Zhou, J.M. (2021). The ZAR1 resistosome is a calcium-permeable channel triggering plant immune signaling. Cell 184, 3528–3541.e3512.

Chang, J.H., Tai, Y.S., Bernal, A.J., Lavelle, D.T., Staskawicz, B.J., and Michelmore, R.W. (2002). Functional analyses of the *Pto* resistance gene family in tomato and the identification of a minor resistance determinant in a susceptible haplotype. Mol Plant Microbe Interact 15, 281–291.

Chen, H., Chen, J., Li, M., Chang, M., Xu, K., Shang, Z., Zhao, Y., Palmer, I., Zhang, Y., McGill, J., Alfano, J.R., Nishimura, M.T., Liu, F., and Fu, Z.Q. (2017). A bacterial type III effector targets the master regulator of salicylic acid signaling, NPR1, to subvert plant immunity. Cell Host Microbe 22, 777–788.e777.

del Pozo, O., Pedley, K.F., and Martin, G.B. (2004). MAPKKKalpha is a positive regulator of cell death associated with both plant immunity and disease. Embo j 23, 3072–3082.

Förderer, A., Li, E., Lawson, A.W., Deng, Y.N., Sun, Y., Logemann, E., Zhang, X., Wen, J., Han, Z., Chang, J., Chen, Y., Schulze-Lefert, P., and Chai, J. (2022). A wheat resistosome defines common principles of immune receptor channels. Nature 610, 532–539.

Franck, C.M., Westermann, J., and Boisson-Dernier, A. (2018). Plant malectin-like receptor kinases: From cell wall integrity to immunity and beyond. Annu Rev Plant Biol 69, 301–328.

Gimenez-Ibanez, S., Hann, D.R., Ntoukakis, V., Petutschnig, E., Lipka, V., and Rathjen, J.P. (2009). AvrPtoB targets the LysM receptor kinase CERK1 to promote bacterial virulence on plants. Curr Biol 19, 423–429.

Göhre, V., Spallek, T., Häweker, H., Mersmann, S., Mentzel, T., Boller, T., de Torres, M., Mansfield, J.W., and Robatzek, S. (2008). Plant pattern-recognition receptor FLS2 is directed for degradation by the bacterial ubiquitin ligase AvrPtoB. Curr Biol 18, 1824–1832.

Gong, Z., and Han, G.Z. (2021). Flourishing in water: the early evolution and diversification of plant receptor-like kinases. Plant J 106, 174–184.

Gong, Z., Qi, J., Hu, M., Bi, G., Zhou, J.M., and Han, G.Z. (2022). The origin and evolution of a plant resistosome. Plant Cell 34, 1600–1620.

Gutierrez, J.R., Balmuth, A.L., Ntoukakis, V., Mucyn, T.S., Gimenez-Ibanez, S., Jones, A.M., and Rathjen, J.P. (2010). Prf immune complexes of tomato are oligomeric and contain multiple Pto-like kinases that diversify effector recognition. Plant J 61, 507–518.

Haruta, M., Sabat, G., Stecker, K., Minkoff, B.B., and Sussman, M.R. (2014). A peptide hormone and its receptor protein kinase regulate plant cell expansion. Science 343, 408–411.

Hoang, D.T., Chernomor, O., von Haeseler, A., Minh, B.Q., and Vinh, L.S. (2018). UFBoot2: Improving the ultrafast bootstrap approximation. Mol Biol Evol 35, 518–522.

Huang, J., Xu, W., Zhai, J., Hu, Y., Guo, J., Zhang, C., Zhao, Y., Zhang, L., Martine, C., Ma, H., and Huang, C.H. (2023). Nuclear phylogeny and insights into whole-genome duplications and reproductive development of Solanaceae plants. Plant Commun 4, 100595.

Janjusevic, R., Abramovitch, R.B., Martin, G.B., and Stebbins, C.E. (2006). A bacterial inhibitor of host programmed cell death defenses is an E3 ubiquitin ligase. Science 311, 222–226.

Ji, D., Liu, W., Cui, X., Liu, K., Liu, Y., Huang, X., Li, B., Qin, G., Chen, T., and Tian, S. (2023). A receptor-like kinase SlFERL mediates immune responses of tomato to *Botrytis cinerea* by recognizing BcPG1 and fine-tuning MAPK signaling. New Phytol 240, 1189–1201.

Jones, J.D., and Dangl, J.L. (2006). The plant immune system. Nature 444, 323–329.

Kalyaanamoorthy, S., Minh, B.Q., Wong, T.K.F., von Haeseler, A., and Jermiin, L.S. (2017). ModelFinder: fast model selection for accurate phylogenetic estimates. Nat Methods 14, 587–589.

Katoh, K., and Standley, D.M. (2013). MAFFT multiple sequence alignment software version 7: improvements in performance and usability. Mol Biol Evol 30, 772–780.

Kim, Y.J., Lin, N.C., and Martin, G.B. (2002). Two distinct *Pseudomonas* effector proteins interact with the Pto kinase and activate plant immunity. Cell 109, 589–598.

Kraus, C.M., Munkvold, K.R., and Martin, G.B. (2016). Natural variation in tomato reveals differences in the recognition of AvrPto and AvrPtoB effectors from *Pseudomonas syringae*. Mol Plant 9, 639–649.

Letunic, I., and Bork, P. (2019). Interactive Tree Of Life (iTOL) v4: recent updates and new developments. Nucleic Acids Res 47, W256–w259.

Li, N., He, Q., Wang, J., Wang, B., Zhao, J., Huang, S., Yang, T., Tang, Y., Yang, S., Aisimutuola, P., Xu, R., Hu, J., Jia, C., Ma, K., Li, Z., Jiang, F., Gao, J., Lan, H., Zhou, Y., Zhang, X., Huang, S., Fei, Z., Wang, H., Li, H., and Yu, Q. (2023). Super-pangenome analyses highlight genomic diversity and structural variation across wild and cultivated tomato species. Nat Genet 55, 852–860.

Lin, N.C., and Martin, G.B. (2005). An avrPto/avrPtoB mutant of *Pseudomonas syringae* pv. *tomato* DC3000 does not elicit Pto-mediated resistance and is less virulent on tomato. Mol Plant Microbe Interact 18, 43–51.

Lin, W., Tang, W., Pan, X., Huang, A., Gao, X., Anderson, C.T., and Yang, Z. (2022). Arabidopsis pavement cell morphogenesis requires FERONIA binding to pectin for activation of ROP GTPase signaling. Curr Biol 32, 497–507.e494.

Liu, C., Shen, L., Xiao, Y., Vyshedsky, D., Peng, C., Sun, X., Liu, Z., Cheng, L., Zhang, H., Han, Z., Chai, J., Wu, H.M., Cheung, A.Y., and Li, C. (2021). Pollen PCP-B peptides unlock a stigma peptide-receptor kinase gating mechanism for pollination. Science 372, 171–175.

Liu, F., Yang, Z., Wang, C., You, Z., Martin, R., Qiao, W., Huang, J., Jacob, P., Dangl, J.L., Carette, J.E., Luan, S., Nogales, E., and Staskawicz, B.J. (2024a). Activation of the helper NRC4 immune receptor forms a hexameric resistosome. Cell 187, 4877–4889.e4815.

Liu, M.J., Yeh, F.J., Yvon, R., Simpson, K., Jordan, S., Chambers, J., Wu, H.M., and Cheung, A.Y. (2024b). Extracellular pectin-RALF phase separation mediates FERONIA global signaling function. Cell 187, 312–330.e322.

Liu, P., Zhao, Z., Tang, Y., Zhou, Y., Liu, J., Xu, K., Chen, Y., Li, X., Tang, Y., and Yang, L. (2025). The HY5-NPR1 module governs light-dependent virulence of a plant bacterial pathogen. Cell Host Microbe 33, 1606–1622.e1610.

Liu, Y., Mahmud, M.R., Xu, N., and Liu, J. (2022). The *Pseudomonas syringae* effector AvrPtoB targets abscisic acid signaling pathway to promote its virulence in Arabidopsis. Phytopathology Research 4, 1–12.

Lu, S., Wang, J., Chitsaz, F., Derbyshire, M.K., Geer, R.C., Gonzales, N.R., Gwadz, M., Hurwitz, D.I., Marchler, G.H., Song, J.S., Thanki, N., Yamashita, R.A., Yang, M., Zhang, D., Zheng, C., Lanczycki, C.J., and Marchler-Bauer, A. (2020). CDD/SPARCLE: the conserved domain database in 2020. Nucleic Acids Res 48, D265–d268.

Luo, X., Xu, N., Huang, J., Gao, F., Zou, H., Boudsocq, M., Coaker, G., and Liu, J. (2017). A lectin receptor-like kinase mediates pattern-triggered salicylic acid signaling. Plant Physiol 174, 2501–2514.

Ma, S., An, C., Lawson, A.W., Cao, Y., Sun, Y., Tan, E.Y.J., Pan, J., Jirschitzka, J., Kümmel, F., Mukhi, N., Han, Z., Feng, S., Wu, B., Schulze-Lefert, P., and Chai, J. (2024). Oligomerization-mediated autoinhibition and cofactor binding of a plant NLR. Nature 632, 869–876.

Madhuprakash, J., Toghani, A., Contreras, M.P., Posbeyikian, A., Richardson, J., Kourelis, J., Bozkurt, T.O., Webster, M.W., and Kamoun, S. (2024). A disease resistance protein triggers oligomerization of its NLR helper into a hexameric resistosome to mediate innate immunity. Sci Adv 10, eadr2594.

Martin, G.B., Frary, A., Wu, T., Brommonschenkel, S., Chunwongse, J., Earle, E.D., and Tanksley, S.D. (1994). A member of the tomato *Pto* gene family confers sensitivity to fenthion resulting in rapid cell death. Plant Cell 6, 1543–1552.

Martin, G.B., Brommonschenkel, S.H., Chunwongse, J., Frary, A., Ganal, M.W., Spivey, R., Wu, T., Earle, E.D., and Tanksley, S.D. (1993). Map-based cloning of a protein kinase gene conferring disease resistance in tomato. Science 262, 1432–1436.

Mucyn, T.S., Clemente, A., Andriotis, V.M., Balmuth, A.L., Oldroyd, G.E., Staskawicz, B.J., and Rathjen, J.P. (2006). The tomato NBARC-LRR protein Prf interacts with Pto kinase in vivo to regulate specific plant immunity. Plant Cell 18, 2792–2806.

Ngou, B.P.M., Ahn, H.K., Ding, P., and Jones, J.D.G. (2021). Mutual potentiation of plant immunity by cell-surface and intracellular receptors. Nature 592, 110–115.

Nguyen, L.T., Schmidt, H.A., von Haeseler, A., and Minh, B.Q. (2015). IQ-TREE: a fast and effective stochastic algorithm for estimating maximum-likelihood phylogenies. Mol Biol Evol 32, 268–274.

Nicholas, K.B. (1997). GeneDoc: Analysis and visualization of genetic variation, EMBNEW. Embnew News 4.

Ntoukakis, V., Balmuth, A.L., Mucyn, T.S., Gutierrez, J.R., Jones, A.M., and Rathjen, J.P. (2013). The tomato Prf complex is a molecular trap for bacterial effectors based on Pto transphosphorylation. PLoS Pathog 9, e1003123.

Ntoukakis, V., Mucyn, T.S., Gimenez-Ibanez, S., Chapman, H.C., Gutierrez, J.R., Balmuth, A.L., Jones, A.M., and Rathjen, J.P. (2009). Host inhibition of a bacterial virulence effector triggers immunity to infection. Science 324, 784–787.

Pedley, K.F., and Martin, G.B. (2004). Identification of MAPKs and their possible MAPK kinase activators involved in the Pto-mediated defense response of tomato. J Biol Chem 279, 49229–49235.

Pilotti, M., Brunetti, A., Uva, P., Lumia, V., Tizzani, L., Gervasi, F., Iacono, M., and Pindo, M. (2014). Kinase domain-targeted isolation of defense-related receptor-like kinases (RLK/Pelle) in Platanus×acerifolia: phylogenetic and structural analysis. BMC Res Notes 7, 884.

Price, M.N., Dehal, P.S., and Arkin, A.P. (2010). FastTree 2–approximately maximum-likelihood trees for large alignments. PLoS One 5, e9490.

Rose, L.E., Langley, C.H., Bernal, A.J., and Michelmore, R.W. (2005). Natural variation in the *Pto* pathogen resistance gene within species of wild tomato (Lycopersicon). I. Functional analysis of Pto alleles. Genetics 171, 345–357.

Rosebrock, T.R., Zeng, L., Brady, J.J., Abramovitch, R.B., Xiao, F., and Martin, G.B. (2007). A bacterial E3 ubiquitin ligase targets a host protein kinase to disrupt plant immunity. Nature 448, 370–374.

Salmeron, J.M., Oldroyd, G.E., Rommens, C.M., Scofield, S.R., Kim, H.S., Lavelle, D.T., Dahlbeck, D., and Staskawicz, B.J. (1996). Tomato Prf is a member of the leucine-rich repeat class of plant disease resistance genes and lies embedded within the *Pto* kinase gene cluster. Cell 86, 123–133.

Saur, I.M.-L., Conlan, B.F., and Rathjen, J.P. (2015). The N-terminal domain of the tomato immune protein Prf contains multiple homotypic and Pto kinase interaction sites. Journal of Biological Chemistry 290, 11258–11267.

Selvaraj, M., Toghani, A., Pai, H., Sugihara, Y., Kourelis, J., Yuen, E.L.H., Ibrahim, T., Zhao, H., Xie, R., Maqbool, A., De la Concepcion, J.C., Banfield, M.J., Derevnina, L., Petre, B., Lawson, D.M., Bozkurt, T.O., Wu, C.H., Kamoun, S., and Contreras, M.P. (2024). Activation of plant immunity through conversion of a helper NLR homodimer into a resistosome. PLoS Biol 22, e3002868.

Sheikh, A.H., Zacharia, I., Pardal, A.J., Dominguez-Ferreras, A., Sueldo, D.J., Kim, J.G., Balmuth, A., Gutierrez, J.R., Conlan, B.F., Ullah, N., Nippe, O.M., Girija, A.M., Wu, C.H., Sessa, G., Jones, A.M.E., Grant, M.R., Gifford, M.L., Mudgett, M.B., Rathjen, J.P., and Ntoukakis, V. (2023). Dynamic changes of the Prf/Pto tomato resistance complex following effector recognition. Nat Commun 14, 2568.

Stegmann, M., Monaghan, J., Smakowska-Luzan, E., Rovenich, H., Lehner, A., Holton, N., Belkhadir, Y., and Zipfel, C. (2017). The receptor kinase FER is a RALF-regulated scaffold controlling plant immune signaling. Science 355, 287–289.

Venkatesh, J., Jahn, M., and Kang, B.C. (2016). Genome-wide analysis and evolution of the Pto-like protein kinase (PLPK) gene family in pepper. PLoS One 11, e0161545.

Wang, J., Hu, M., Wang, J., Qi, J., Han, Z., Wang, G., Qi, Y., Wang, H.W., Zhou, J.M., and Chai, J. (2019). Reconstitution and structure of a plant NLR resistosome conferring immunity. Science 364.

Wang, M.Y., Chen, J.B., Wu, R., Guo, H.L., Chen, Y., Li, Z.J., Wei, L.Y., Liu, C., He, S.F., Du, M.D., Guo, Y.L., Peng, Y.L., Jones, J.D.G., Weigel, D., Huang, J.H., and Zhu, W.S. (2023). The plant immune receptor SNC1 monitors helper NLRs targeted by a bacterial effector. Cell Host Microbe 31, 1792–1803.e1797.

Wang, Y., Zhang, Y., Gao, Z., and Yang, W. (2018). Breeding for resistance to tomato bacterial diseases in China: Challenges and prospects. Horticultural Plant Journal 4, 193–207.

Wei, H.L., Chakravarthy, S., Worley, J.N., and Collmer, A. (2013). Consequences of flagellin export through the type III secretion system of *Pseudomonas syringae* reveal a major difference in the innate immune systems of mammals and the model plant *Nicotiana benthamiana*. Cell Microbiol 15, 601–618.

Xin, X.F., and He, S.Y. (2013). *Pseudomonas syringae* pv. *tomato* DC3000: a model pathogen for probing disease susceptibility and hormone signaling in plants. Annu Rev Phytopathol 51, 473–498.

Xu, F., Wang, L., Li, Y., Shi, J., Staiger, D., and Yu, F. (2024). Phase separation of GRP7 facilitated by FERONIA-mediated phosphorylation inhibits mRNA translation to modulate plant temperature resilience. Mol Plant 17, 460–477.

Xu, N., Luo, X., Wu, W., Xing, Y., Liang, Y., Liu, Y., Zou, H., Wei, H.L., and Liu, J. (2020). A plant lectin receptor-like kinase phosphorylates the bacterial effector AvrPtoB to dampen its virulence in Arabidopsis. Mol Plant 13, 1499–1512.

Yuan, M., Jiang, Z., Bi, G., Nomura, K., Liu, M., Wang, Y., Cai, B., Zhou, J.M., He, S.Y., and Xin, X.F. (2021). Pattern-recognition receptors are required for NLR-mediated plant immunity. Nature 592, 105–109.

Zhang, N., Gan, J., Carneal, L., González-Tobón, J., Filiatrault, M., and Martin, G.B. (2024). Helper NLRs Nrc2 and Nrc3 act codependently with Prf/Pto and activate MAPK signaling to induce immunity in tomato. Plant J 117, 7–22.

Zhang, X., Yang, Z., Wu, D., and Yu, F. (2020). RALF–FERONIA signaling: Linking plant immune response with cell growth. Plant Communications 1, 100084.

