## Supplemental Figures for "Pto evolved from malectin-like RLKs phosphorylates AvrPtoB to promote Prf-mediated immunity in *Solanum pimpinellifolium*"

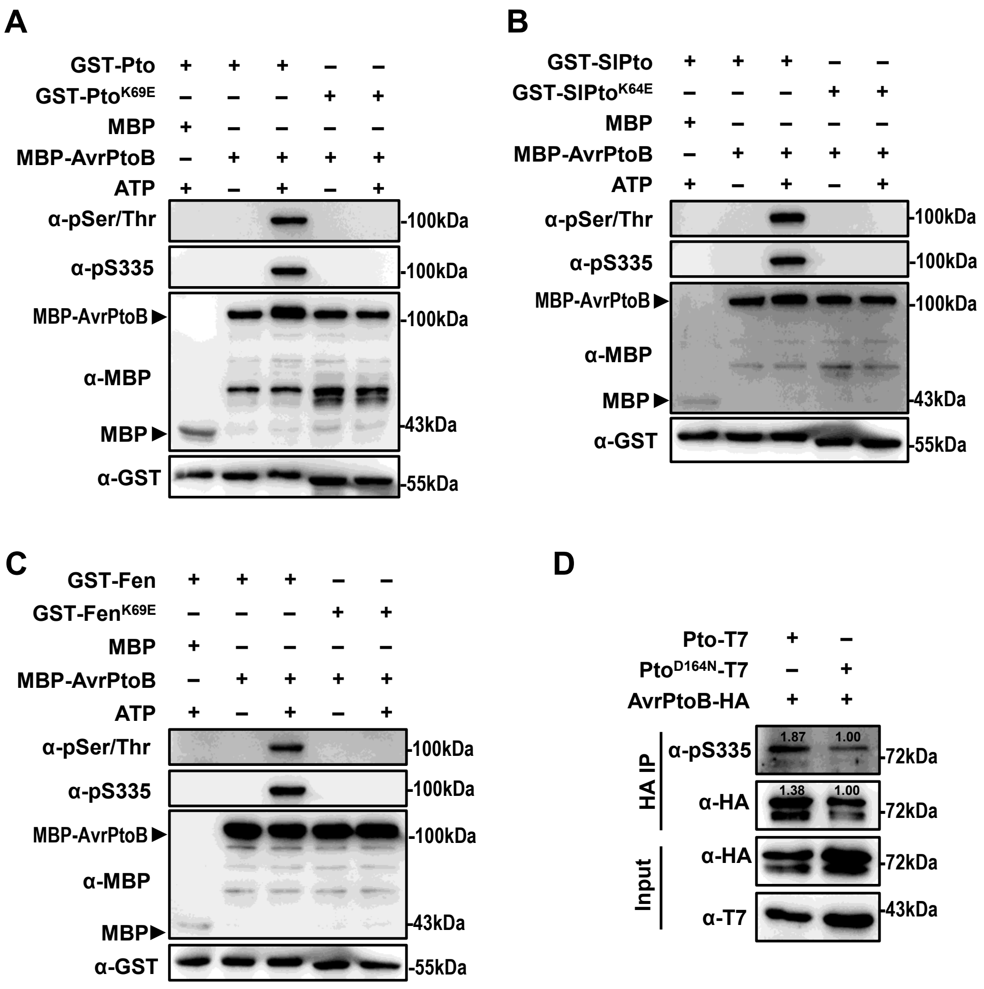


**Supplementary Figure S1. Kinase-dead mutants of Pto, SlPto, and Fen can't phosphorylate AvrPtoB at Ser335 (Supports figure 1). A** to **C)** Recombinant GST-Pto (from *S. pimpinellifolium*), GST-SlPto (from *S. lycopersicum*), GST-Fen (from *S. pimpinellifolium*), MBP-AvrPtoB, and kinase-dead mutants were subjected to kinase assay. Protein phosphorylation was identified by immunoblot using anti-phosphoserine/threonine antibody (α-pSer/Thr). The phosphorylated AvrPtoB was detected by anti-pS335 antibody (α-pS335). **D)** Pto phosphorylates AvrPtoB at Ser335 site in *N. benthamiana*. *35S:Pto-T7*, *35S:Pto$^{D164N}$-T7* was co-expressed with *35S:AvrPtoB-HA* in *N. benthamiana*. The samples were collected at 36 hpi for HA-IP assay. The phosphorylation of AvrPtoB at Ser335 site was detected by α-pS335.

```
S. lycopersicoides   MGSKYSKATNSTSDASNSRSGVPFESYRVPFVDLEEATNNFDEKFFIAEGGFGKVYRGVLRDGTKVALKRHNRDSGQSIKEFRTEIEILSRCSHPHLVSLIGYCDER 107
S. habrochaites      MGSKYSKATNSINDALSSSYLVPFESYRVPLVDLEEATNNFDDKFFIGAGVFGKVYKGVLRDGTKVALKRRKPESSQGIEELETEIEILSFCRHPHLVSLIGFCDER 107
S. pennellii         MGSKYSKATNSINDALSSSYLVPFESYRVPLVDLEEATNNFDDKFFIGAGVFGKVYKGVLRDGTKVALKRRKPESSQGIEELETEIEILSFCRHPHLVSLIGFCDER 107
S. chilense          MGSKYSKATNSINDALSSSYLVPFESYRVPLVDLEEATNNFDDKFLIGHGLFGKVYKGVLRDGTKVALKRRQPESSQGIEEFETEIETLSFCRHPHLVSLIGFCDER 107
S. peruvianum        MGSKYSKATNSINDALSSSYLVPFESYRVPLVDLEEATNNFDHKFLTGAGVFGKVYKGVLRDGTKVALKRRQPESSQGIEEFETEIETLSFCRHPHLVSLIGFCDER 107
S. corneliomulleri   MGSKYSKATNSINDALSSSYLVPFESYRVPLVDLEEATNNFDDKFFIGEGGFGKVYKGVLRDGTKVALKSCKPESSQGIEEFETEIEILSFCRHPHLVSLIGFCDER 107
S. neorickii         MGSKYSKATNSISDASNSRYGVPFENYQVPLVDLEEATNNFDDNFFIAEGGFGKVYRGVLRDGTKVALKRHNCDSQQSIEEFRTEIDILSRRSHPHLVSLIGYCDGR 107
S. chmielewskii      MGSKYSKATNSINDALSSSYLVPFESYRVPLVDLEEATNNFDDKFFIGEGGFGKVYKGVLRDGTKVALKSCKPESSQGIEEFETEIEILSFCRHPHLVSLIGFCDER 107
S. pimpinellifolium  MGSKYSKATNSINDALSSSYLVPFESYRVPLVDLEEATNNFDHKFLIGHGVFGKVYKGVLRDGAKVALKRRTPESSQGIEEFETEIETLSFCRHPHLVSLIGFCDER 107
S. galapagense       MGSKYSKATNSISDASNS-----FESYRFPLEDLEEATKNFDDKFFIGEGAFGKVYKGVLRDGTKVALKRQNRDSRQGIEEFGTEIGILSRRSHPHLVSLIGYCDER 102
S. lycopersicum      MGSKYSKATNSISDASNS-----FESYRFPLEDLEEATNNFDDKFFIGEGAFGKVYKGVLRDGTKVALKRQNRDSRQGIEEFGTEIGILSRRSHPHLVSLIGYCDER 102

S. lycopersicoides   NEMIVVYDYIENGNLKSHLYGSDLP--TMTWEQRLEICIGAARALHYLHTNGLIHRDVKSINILLDENFVPKITDFGLSKIRPELDQTHVSTVVKGTFGYLDPEYYH 212
S. habrochaites      NEMILIYKYMENGNLKRHLYGSDLP--SMSWEQRLEICIGAARGLHFIHTRAVIHRDVKSINILLDENFVPKITDFGLSKKGTELDQTHLSTVVQGTLGYLDPEYFI 212
S. pennellii         NEMILIYKYMENGNLKRHLYGSDLP--SMSWEQRLEICIGAARGLHFIHTRAVIHRDVKSINILLDENFVPKITDFGLSKKGTELDQTHLSTVVQGTLGYLDPEYFI 212
S. chilense          NEMILIYKYMENGNLKRHLYGSDLP--SMSWEQRLEICIGAARGLHYLHTRAVIHRDVKSINILLDENFVPKITDFGISKKGTELDQTHLSTVVKGTLGYLDPEYFI 212
S. peruvianum        NEMILIYKYMENGNLNRHLCGSDLP--SMSWEQRLEICIGAARGLHYLHTRAVIHRDVKSTNILLDENFVAKITDFGISKKGTELDQTHLSTDVKGTFGYLDPEYFI 212
S. corneliomulleri   NEMILIYKYMENGNLKRHLYGSDLL--FMSWEQRLEICIGAARGLHYLHTRAIIHRDVKSINILLDENFVAKITDFGISKKGTELDQTHVSTVVKGTLGYLDPEYFI 212
S. neorickii         NEMILIYDYMENGNLKSHLYGSDLP--SMSWEQRLEICIGAARGLHYLHTNGVMHRDVKSSNILLDENFVPKITDFGLSKTRPQLYQTHVSTDVKGTFGYIDPEYFI 212
S. chmielewskii      NEMILIYKYMENGNLKRHLYGSDLP--SMSWEQRLEICIGAARGLHYLHTRAIIHRDVKSINILLDENFVPKITDFGISKKGTELDQTHLSTVVKGTLGYLDPEYFI 212
S. pimpinellifolium  NEMILIYKYMENGNLKRHLYGSDLPTMSMSWEQRLEICIGAARGLHYLHTRAIIHRDVKSINILLDENFVPKITDFGISKKGTELDQTHLSTVVKGTLGYIDPEYFI 214
S. galapagense       NEMVLIYDYMENGNLKSHLTGSDLP--SMSWEQRLEICIGAARGLHYLHTNGVMHRDVKSSNILLDENFVPKITDFGLSKTRPQLYQT---TDVKGTFGYIDPEYFI 204
S. lycopersicum      NEMVLIYDYMENGNLKSHLTGSDLP--SMSWEQRLEICIGAARGLHYLHTNGVMHRDVKSSNILLDENFVPKITDFGLSKTRPQLYQT---TDVKGTFGYIDPEYFI 204

S. lycopersicoides   RGQLTEKSDVYSFGVVLLEVLCARSAIVQSLPRKMVSLAGWAVESHNNGQLEQIIDRNLVAKIRPESLRKFGETAVKCLALSGKDRPSMGDVLWKLEYALRLQESVI 319
S. habrochaites      KGRLTEKSDVYSFGVVLFEVLCARSAIVQSLPREMVNLAEWAVESHNNGQLEQIIDPNLADKIRPESLRKFGETAVKCLALSSEDRPSMGDVLWKLEYALRLQESVI 319
S. pennellii         KGRLTEKSDVYSFGVVLFEVLCARSAIVQSLPREMVNLAEWAVESHNNGQLEQIIDPNLADKIRPESLRKFGETAVKCLALSSEDRPSMGDVLWKLEYALRLQESVI 319
S. chilense          KGRLTEKSDVYSFGVVLFEVLCARSAIVQSLPREMVNLAEWAVESHNNGQLEQIVDPNLADKIRPESLRKFGETAVKCLALSSEDRPSMGDVLWKLEYALRLQESVI 319
S. peruvianum        KGRLTEKSDVYSFGVVLFEVLCARSTIVQSLPREMVNLAEWAVESHNNGQLEQIVDPNIADKIRPESLRKFGETAVKCLALSSEDRPSMGDVLWKLEYALRLQESVI 319
S. corneliomulleri   KGRLTEKSDVYSFGVVLFEVLCARSAIVQSLPREMVNLAEWAVESHNNGQLEQIVDPNIADKIRPESLRKFGETAVKCLALSSEDRPSMGDVLWKLEYALRLQESVI 319
S. neorickii         KGRLTEKSDVYSFGVVLFEVLCARSAIVQSLPSEMVNLAEWAVESHNNGQLEQIIDPNLAAKIRPESLRKFGETAVKCLALSSEDRPSMGDVLWKLEYALRLQESVI 319
S. chmielewskii      KGRLTEKSDVYSFGVVLFEVLCARSAIVQSLPREMVNLAEWAVESHNNGQLEQIVDPNLADKIRPESLRKFGETAVKCLALSSEDRPSMGDVLWKLEYALRLQESVI 319
S. pimpinellifolium  KGRLTEKSDVYSFGVVLFEVLCARSAIVQSLPREMVNLAEWAVESHNNGQLEQIVDPNLADKIRPESLRKFGDTAVKCLALSSEDRPSMGDVLWKLEYALRLQESVI 321
S. galapagense       KGRLTEKSDVYSFGVVLFEVLCARSAMVQSLPREMVNLAEWAVESHNNGQLEQIVDPNLADKIRPESLRKFGETAVKCLALSSEDRPSMGDVLWKLEYALRLQESVI 311
S. lycopersicum      KGRLTEKSDVYSFGVVLFEVLCARSAMVQSLPREMVNLAEWAVESHNNGQLEQIVDPNLADKIRPESLRKFGETAVKCLALSSEDRPSMGDVLWKLEYALRLQESVI 311
```

**Supplementary Figure S2. Multiple sequence alignment of Pto and its homologous proteins from wild and cultivated tomato species (Supports figure 1).** Sequence alignment was performed using Clustal W (https://www.genome.jp/tools-bin/clustalw). Red, blue and green backgrounds represented the amino acid similarity reaching 100%, 80% and 60%, respectively. Kinase domain (amino acids from 47 to 311) of Pto was marked with red line. The activation domain (amino acid from 182-214) of Pto was marked with the green line. The P+1 loop (amino acids from 201 to 210) of Pto was marked with blue line. ATP binding site (Lys-69) of Pto was marked with the red box.

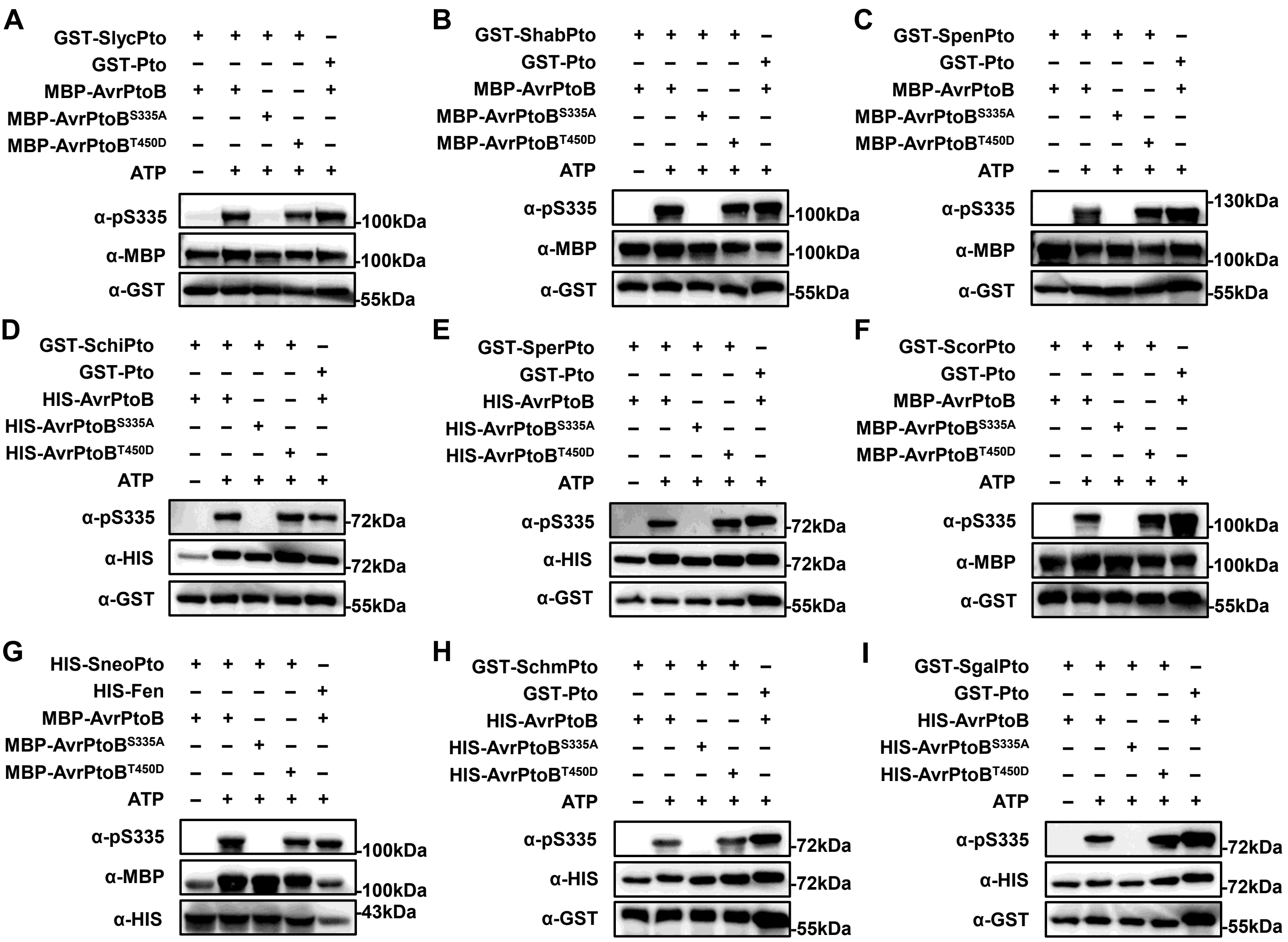


**Supplementary Figure S3. Pto homologs phosphorylates AvrPtoB at Ser335 (Supports figure 1). A to I)** Recombinant GST-Pto, HIS-Fen, GST-Pto-homologs, HIS-SneoPto， HIS-AvrPtoB, MBP-AvrPtoB and site-mutated variants AvrPtoB$^{S335A}$ and AvrPtoB$^{T450D}$ were subjected to kinase activity detection assay. The phosphorylated AvrPtoB was detected by anti-pS335 antibody (α-pS335). The sequences of *SlycPto*, *ShabPto*, *SpenPto*, *SchiPto*, *SperPto*, *ScorPto*, *SneoPto*, *SchmPto* and *SgalPto* were cloned from *S. lycopersicoides*, *S. habrochaites*, *S. pennellii*, *S. chilense*, *S. peruvianum*, *S. corneliomulleri*, *S. neorickii*, *S. chmielewskii* and *S. galapagense*, respectively. GST-Pto or HIS-Fen was used as positive control.

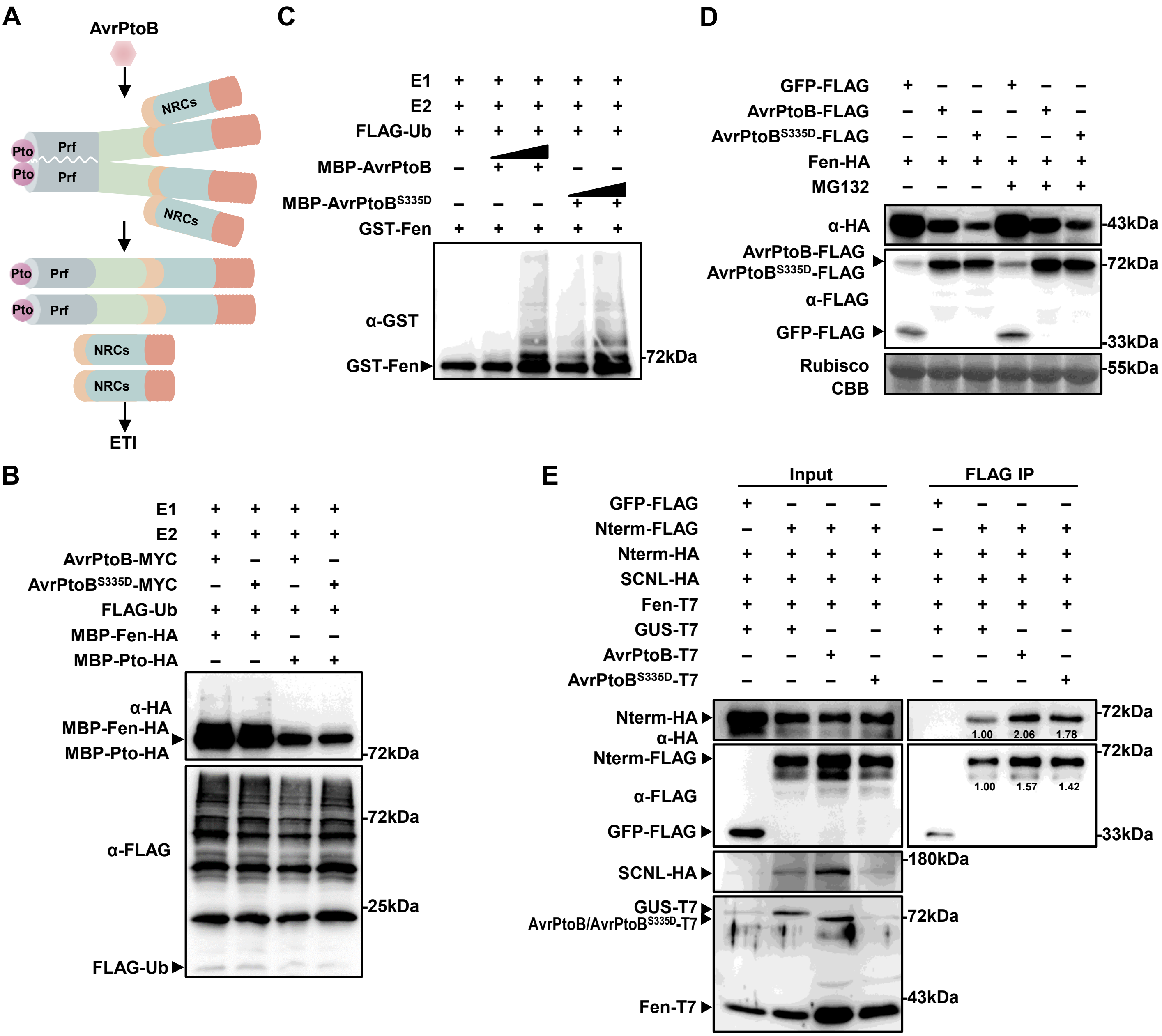


**Supplementary Figure S4. AvrPtoB$^{S335D}$ doesn't dissociate Prf complex through Fen (Supports figure 3). A)** The schematic diagram of Pto/Prf complex recognizing AvrPtoB to trigger ETI response. **B and C)** AvrPtoB and AvrPtoB$^{S335D}$ ubiquitinate Fen *in vitro*. The bacterial lysates from *E. coli* BL21 strains expressing E1 (AtUBA1), E2 (AtUBC8), AvrPtoB-MYC or AvrPtoB$^{S335D}$-MYC, MBP-Fen-HA and FLAG-Ub were detected by immunoblot using anti-HA and anti-FLAG antibody. Pto was used as negative control (**B**). The recombinant proteins were purified from *E.coli* BL21. The triangles showed that the content of MBP-AvrPtoB or MBP-AvrPtoB$^{S335D}$ increased from 0.1 µg to 0.5 µg. The ubiquitinated Fen was detected by anti-GST. E1: wheat; E2: Human UbcH5b (**C**). **D)** AvrPtoB and AvrPtoB$^{S335D}$ degrade Fen through 26S proteasome. *35S: Fen-HA* was coexpressed with *35S: GFP-FLAG*, *35S: AvrPtoB-FLAG* or *35S: AvrPtoB$^{S335D}$-FLAG* in *N. benthamiana*. 100 µM MG132 was used to inhibit 26S proteasome-mediated protein degradation at 24 hpi. Samples were collected at 6-8 h after MG132 treatment. The proteins were detected by immunoblot using corresponding antibody. **E)** AvrPtoB$^{S335D}$ doesn't dissociate Prf complex through Fen by Co-IP assay. *35S:Prf-Nterm-FLAG*, *35S:Prf-Nterm-HA*, *35S:Prf-SCNL-HA*, *35S:Fen-T7* and *35S:AvrPtoB* or *35S:AvrPtoBS$^{335D}$* were transiently expressed in *N. benthamiana*. The immunoprecipitation was performed after 36 hpi. The band intensity was calculated by Image J. The smaller value represents the weaker interaction between proteins. All experiments were repeated three times with similar results.

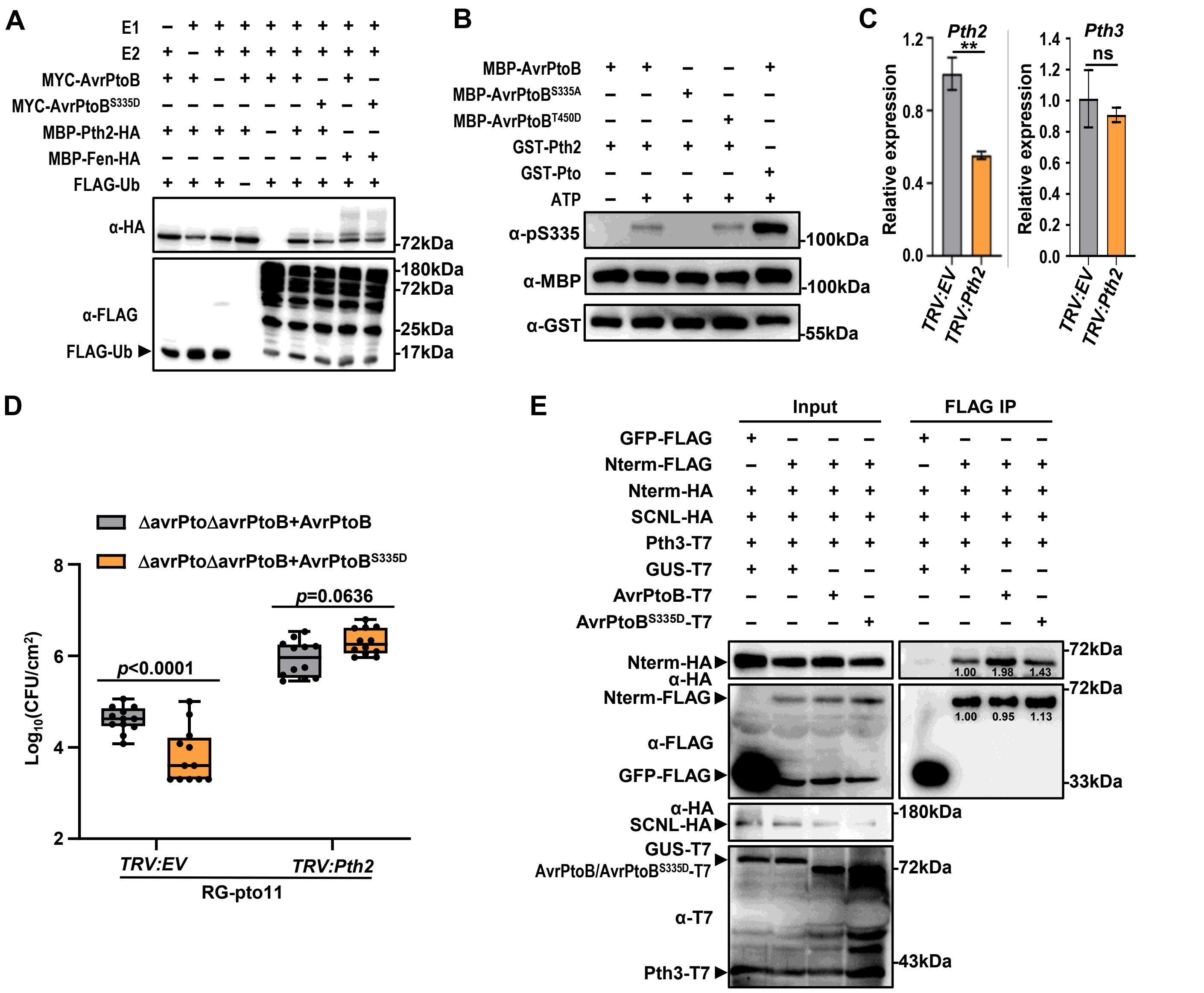


**Supplementary Figure S5. AvrPtoB$^{S335D}$ induces the dissociation of Prf complex through Pth2 but not Pth3 (Supports figure 4). A)** AvrPtoB and AvrPtoB$^{S335D}$ can't ubiquitinate Pth2 *in vitro*. The bacterial lysates from *E. coli* BL21 strains expressing E1 (AtUBA1), E2 (AtUBC8), AvrPtoB-MYC, AvrPtoB$^{S335D}$-MYC, MBP-Pth2-HA or MBP-Fen-HA and FLAG-Ub, or lacking one of these proteins were detected by immunoblot using anti-HA and anti-FLAG antibody. The ubiquitination of Fen by AvrPtoB and AvrPtoB$^{S335D}$ were used as positive control. **B)** Pth2 can weakly phosphorylate AvrPtoB at Ser335 site *in vitro*. GST-Pth2, GST-Pto, MBP-AvrPtoB, MBP-AvrPtoB$^{S335A}$, and MBP-AvrPtoB$^{T450D}$ were subjected to *in vitro* kinase assay. The phosphorylation of AvrPtoB was detected by immunoblot using α-pS/T and α-pS335 antibody. **C)** The expression of *Pth2* and *Pth3* in *Pth2*-scilenced RG-pto11 plants was detected by RT-qPCR. Values are means±SD (n = 3; Student's *t*-test; ***p*< 0.01; ns: no significance). **D)** The silence of *Pth2* weakened the reduced pathogenicity caused by AvrPtoB$^{S335D}$. *Pth2* in RG-pto11 plants was silenced by VIGS. 3 weeks later, tomato plants were inoculated with *Pst* DC3000 ΔavrPtoΔavrPtoB expressing *AvrPtoB* and *AvrPtoB$^{S335D}$* at the concentration of $5\times10^4$ CFU/mL. The plants were subjected to growth curve analysis at 3 dpi. Error bars represent means ± SD (n = 12; two-way ANOVA). *p*-values were marked on the bar. **E)** AvrPtoB$^{S335D}$ can't induce the dissociation of Prf complex through Pth3. *35S: GFP-FLAG*, *35S: Nterm-FLAG*, *35S: Nterm-HA*, *35S: SCNL-HA*, *35S: GUS-T7*, *35S: AvrPtoB-T7*, and *35S: AvrPtoB$^{S335D}$-T7* were co-expressed in *N. benthamiana*. Extracted proteins were immunoprecipitated 36 hpi using anti-FLAG beads. The Co-IP proteins were detected by anti-HA antibody. The band intensity was calculated by Image J. All experiments were repeated three times with similar results.

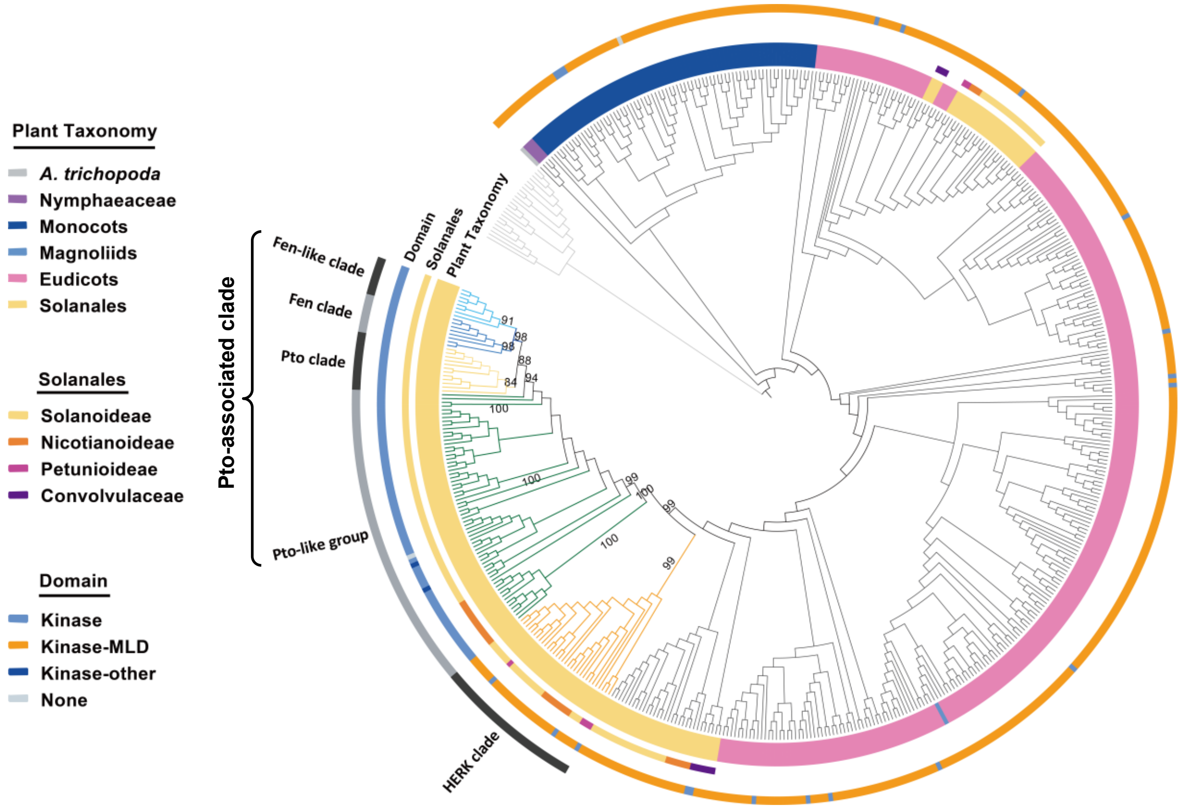


**Supplementary Figure S6. Phylogenetic analysis of the tomato Pto-related proteins (Supports figure 6).** The phylogeny was constructed using the maximum likelihood method. Outgroups were indicated by light gray branches. Different groups were indicated by outermost strip as well as different colors of branches (Fen-like in light blue, Fen in blue, Pto in gold, Pto-like in green and HERK in orange). For each protein, the domain architecture was indicated by the second strip, while the third and the innermost strip indicated the subfamilies of the Solanales and overall classification, respectively. MLD: Malectin-Like Domain. The values near the nodes were UFBoot values.

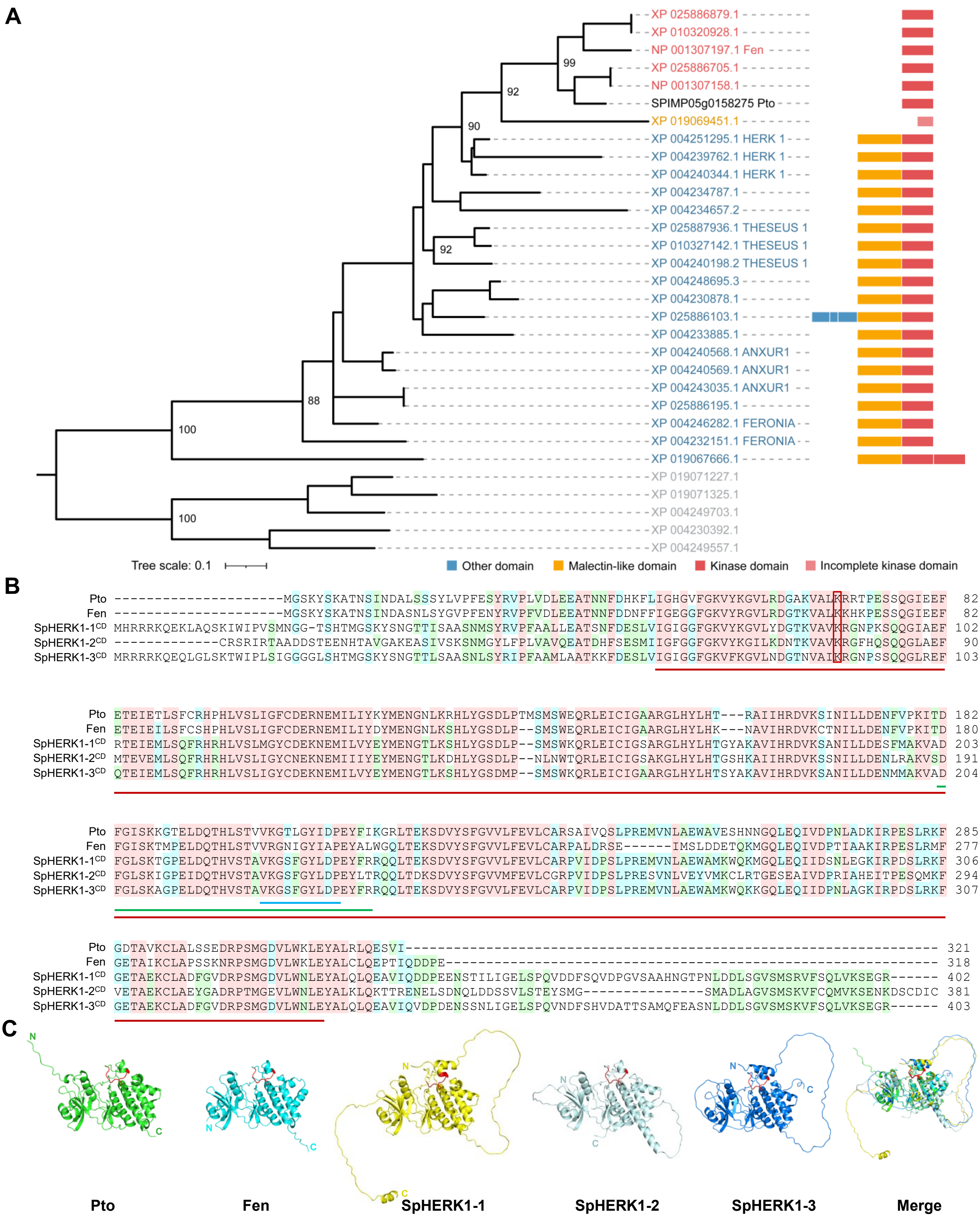
A
XP 025886879.1
XP 010320928.1
NP 001307197.1 Fen
XP 025886705.1
NP 001307158.1
SPIMP05g0158275 Pto
XP 019069451.1
XP 004251295.1 HERK 1
XP 004239762.1 HERK 1
XP 004240344.1 HERK 1
XP 004234787.1
XP 004234657.2
XP 025887936.1 THESEUS 1
XP 010327142.1 THESEUS 1
XP 004240198.2 THESEUS 1
XP 004248695.3
XP 004230878.1
XP 025886103.1
XP 004233885.1
XP 004240568.1 ANXUR1
XP 004240569.1 ANXUR1
XP 004243035.1 ANXUR1
XP 025886195.1
XP 004246282.1 FERONIA
XP 004232151.1 FERONIA
XP 019067666.1
XP 019071227.1
XP 019071325.1
XP 004249703.1
XP 004230392.1
XP 004249557.1
99
92
90
92
88
100
100
Tree scale: 0.1
Other domain
Malectin-like domain
Kinase domain
Incomplete kinase domain
B
Pto ---------------------------MGSKYSKATNSINDALSSSYLVPFESYRVPLVDLEEATNNFDHKFLIGHGVFGKVYKGVLRDGAKVALKRRTPESSQGIEEF 82
Fen ---------------------------MGSKYSKATNSINDASNLSYGVPFENYRVPFVDLEEATNNFDDNFFIGEGGFGKVYRGVLRDGTKVALKKHKPESSQGIEEF 82
SpHERK1-1CD MHRRRKQEKLAQSKIWIPVSMNGG-TSHTMGSKYSNGTTISAASNMSYRVPFAALLEATSNFDESLVIGIGGFGKVYKGVLYDGTKVAVKRGNPKSQQGIAEF 102
SpHERK1-2CD --------------CRSRIRTAADDSTEENHTAVGAKEASIVSKSNMGYLFPLVAVQEATDHFSESMIIGFGGFGKVYKGILKDNTKVAVKRGFHQSQQGLAEF 90
SpHERK1-3CD MRRRRKQEQLGLSKTWIPLSIGGGGLSHTMGSKYSNGTTLSAASNLSYRIPFAAMLAATKKFDESLVIGIGGFGKVFKGVLNDGTNVAIKRGNPSSQQGLREF 103
Pto ETEIETLSFCRHPHLVSLIGFCDERNEMILIYKYMENGNLKRHLYGSDLPTMSMSWEQRLEICIGAARGLHYLHT---RAIIHRDVKSINILLDENFVPKITD 182
Fen ETEIEILSFCSHPHLVSLIGFCDERNEMILIYDYMENGNLKSHLYGSDLP--SMSWEQRLEICIGAARGLHYLHK---NAVIHRDVKCTNILLDENFVPKITD 180
SpHERK1-1CD RTEIEMLSQFRHRHLVSLMGYCDEKNEMILVYEYMENGTLKSHLYGSDLP--SMSWKQRLEICIGSARGLHYLHTGYAKAVIHRDVKSANILLDESFMAKVAD 203
SpHERK1-2CD MTEVEMLSQFRHRHLVSLIGYCNEKNEMIIIYEYMENGTLKDHLYGSDLP--NLNWTQRLEICIGSAKGLHYLHTGSHKAIIHRDVKSSNILLDENLRAKVSD 191
SpHERK1-3CD QTEIEMLSQFRHRHLVSLIGYCDEKNEMILVYEYMENGTLKSHLYGSDMP--SMSWKQRLEICIGAARGLHYLHTSYAKAVIHRDVKSANILLDENMMAKVAD 204
Pto FGISKKGTELDQTHLSTVVKGTLGYIDPEYFIKGRLTEKSDVYSFGVVLFEVLCARSAIVQSLPREMVNLAEWAVESHNNGQLEQIVDPNLADKIRPESLRKF 285
Fen FGISKTMPELDQTHLSTVVRGNIGYIAPEYALWGQLTEKSDVYSFGVVLFEVLCARPALDRSE------IMSLDDETQKMGQLEQIVDPTIAAKIRPESLRMF 277
SpHERK1-1CD FGLSKTGPELDQTHVSTAVKGSFGYLDPEYFRRQQLTEKSDVYSFGVVLFEVLCARPVIDPSLPREMVNLAEWAMKWQKMGQLEQIIDSNLEGKIRPDSLRKF 306
SpHERK1-2CD FGLSKIGPEIDQTHVSTAVKGSFGYLDPEYLTRQQLTDKSDVYSFGVVMFEVLCGRPVIDPSLPRESVNLVEYVMKCLRTGESEAIVDPRIAHEITPESQMKF 294
SpHERK1-3CD FGLSKAGPELDQTHVSTAVKGSFGYLDPEYFRRQQLTEKSDVYSFGVVLFEVLCARPVIDPSLPREMVNLAEWAMKWQKKGQLEQIIDPNLAGKIRPDSLRKF 307
Pto GDTAVKCLALSSEDRPSMGDVLWKLEYALRLQESVI------------------------------------------------------------------ 321
Fen GETAIKCLAPSSKNRPSMGDVLWKLEYALCLQEPTIQDDPE------------------------------------------------------------- 318
SpHERK1-1CD GETAEKCLADFGVDRPSMGDVLWNLEYALQLQEAVIQDDPEENSTILIGELSPQVDDFSQVDPGVSAAHNGTPNLDDLSGVSMSRVFSQLVKSEGR------ 402
SpHERK1-2CD VETAEKCLAEYGADRPTMGEVLWNLEYALKLQKTTRENELSDNQLDDSSVLSTEYSMG---------------SMADLAGVSMSKVFCQMVKSENKDSCDIC 381
SpHERK1-3CD GETAEKCLADFGVDRPSMGDVLWNLEYALQLQEAVIQVDPDENSSNLIGELSPQVNDFSHVDATTSAMQFEASNLDDLSGVSMSKVFSQLVKSEGR------ 403
C
N
C
Pto
Fen
SpHERK1-1
SpHERK1-2
SpHERK1-3
Merge

**Supplementary Figure S7. Homology analysis of Pto and tomato MLR sequence (Supports figure 7). A)** The phylogeny was constructed using the maximum likelihood method. Outgroups including 5 LRR-RLKs were indicated by gray. Different groups were indicated by different colors (Pto in black, proteins with kinase and malectin-like domains in blue, proteins with only a kinase domain in red and the protein with only an incomplete kinase domain in pink). For each protein, the domain architecture was shown next to the protein name. Different domains were indicated using different color stripes, and the color key was shown on the lower right. The values near the nodes were UFBoot values. **B)** The full-length sequences of Pto and Fen, the intracellular domain sequences of SpHERK1-1, SpHERK1-2 and SpHERK1-3 were used to perform sequence alignment. Sequence alignment was performed using Clustal W (https://www.genome.jp/tools-bin/clustalw). Red, blue and green represented the amino acid similarity reaching 100%, 80% and 60%, respectively. Kinase domain of Pto (amino acids from 47 to 311) was marked with red line. The P+1 loop of Pto (amino acids from 201 to 210) was marked with blue line. The activation domain (amino acid from 182-214) of Pto was marked with the green line. ATP binding site of Pto (Lys-69) was marked with the red box. **C)** The structures of Pto, Fen, and the intracellular domain of SpHERK1-1, SpHERK1-2, and SpHERK1-3 were obtained from UniProt (https://www.uniprot.org/). The merged structure was visualized by PyMol software. The P+1 loop was highlight in red.

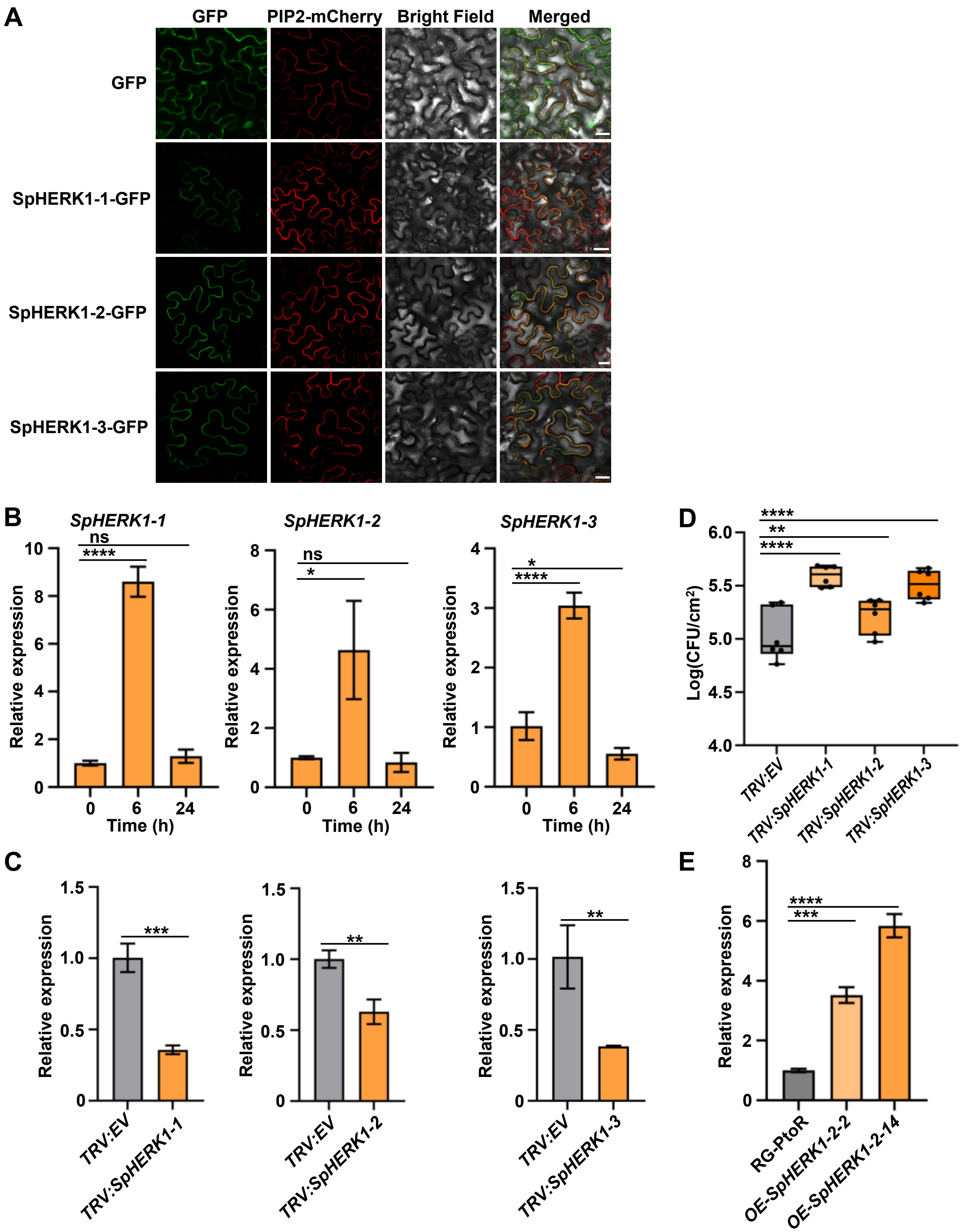


**Supplementary Figure S8. Three SpHERK1 proteins positively regulate tomato disease against *Pst* DC3000 (Supports figure 7). A)** Subcellular localization of SpHERK1-1, SpHERK1-2 and SpHERK1-3. *N. benthamiana* leaves were co-infiltrated with *35S:PIP2-mCherry* and *35S:SpHERK1-1-GFP*, *35S:SpHERK1-2-GFP* or *35S:SpHERK1-3-GFP*. The PIP2-mCherry was used as the plasma membrane marker. The images were obtained by confocal microscope at 36 hpi. Bar = 25 μm. **B)** The expressions of *SpHERK1-1*, *SpHERK1-2* and *SpHERK1-3* are induced by *Pst* DC3000. Four-week-old tomatoes were inoculated with $2\times10^8$ CFU/mL *Pst* DC3000. RT-qPCR was performed at 0, 6, and 24 h. Values are means ± SD (n = 3 biological replicates; Two-way ANOVA; *$p< 0.05$, ****$p< 0.0001$; ns: no significance). **C and D)** Growth curve analysis of *Pst* DC3000 infection in *SpHERK1s* silenced tomato plants. The transcription of *SpHERK1-1*, *SpHERK1-2* and *SpHERK1-3* were knocked down by VIGS in tomato. The expressions of *SpHERK1-1*, *SpHERK1-2* and *SpHERK1-3* in silenced tomatoes were detected by RT-qPCR. Values were means ± SD (n = 3 biological replicates; Student's *t* test; **$p< 0.01$, ***$p< 0.001$) (C). After 3 weeks, silenced tomato plants were inoculated with *Pst* DC3000 at concentration of $2\times10^8$ CFU/mL. Plants were subjected to bacterial growth curve analysis at 3 dpi. Values were means ± SD (n = 6 biological replicates; Two-way ANOVA; **$p< 0.01$, ****$p< 0.0001$) (D). **E)** The expression of *SpHERK1-2* is detected by RT-qPCR in *SpHERK1-2* overexpressed tomato lines. Values are means ± SD (n = 3 biological replicates; Two-way ANOVA; ***$p< 0.001$, ****$p< 0.0001$).

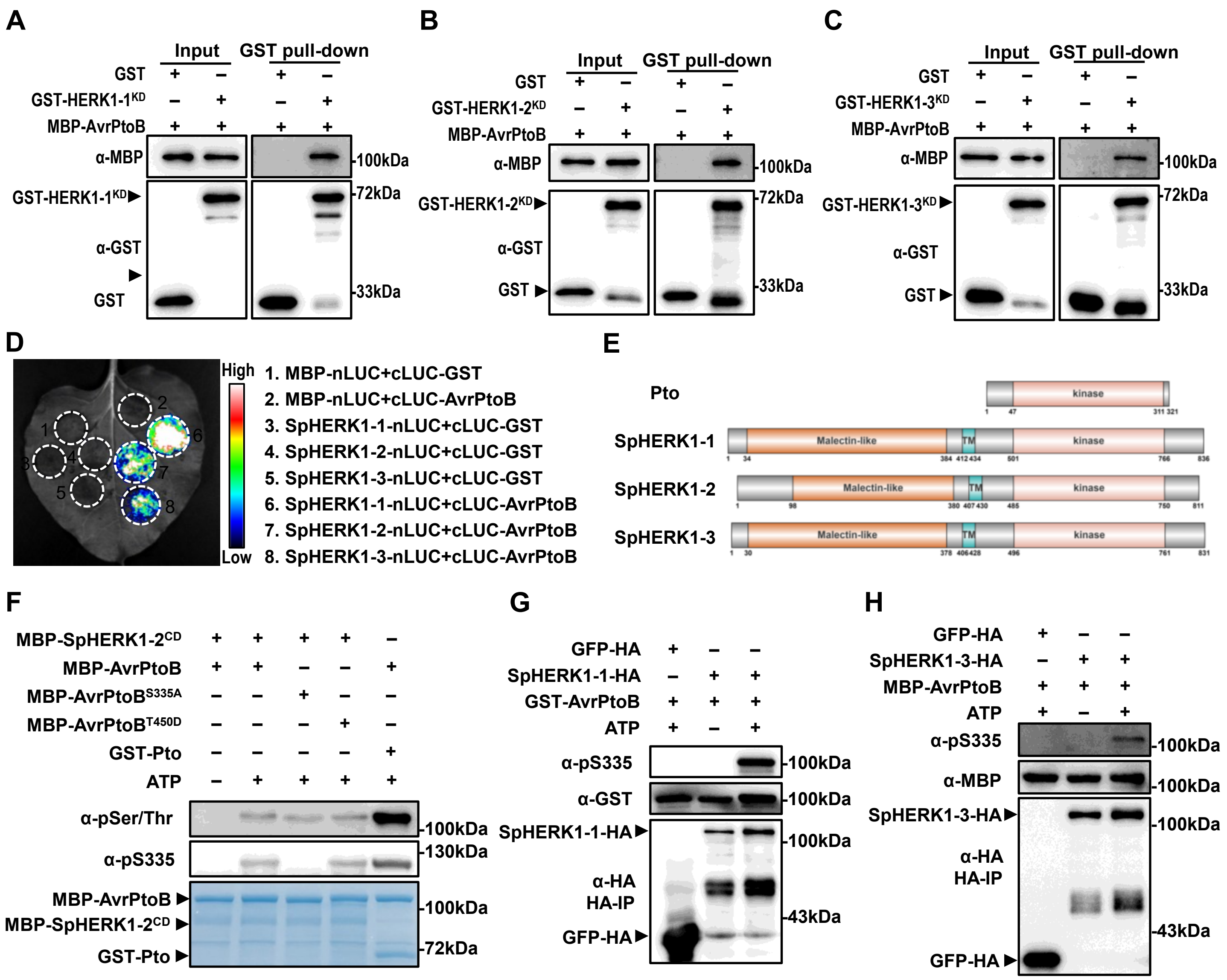


**Supplementary Figure S9. SpHERK1s interact with and phosphorylate AvrPtoB Ser335 site (Supports figure 7). A-C)** AvrPtoB interacts with SpHERK1-1 (**A**), SpHERK1-2 (**B**) and SpHERK1-3 (**C**) *in vitro* by GST pull-down assay. The recombinant MBP-AvrPtoB, GST-HERK1-1$^{KD}$, GST-HERK1-2$^{KD}$, and GST-HERK1-3$^{KD}$ were subjected to GST pull-down assay. The interacting proteins were detected by immunoblot. **D)** AvrPtoB interacts with SpHERK1-1, SpHERK1-2, and SpHERK1-3 *in vivo* by LCA. *35S: cLUC-AvrPtoB* was co-expressed with *35S: SpHERK1-1-nLUC*, *35S: SpHERK1-2-nLUC* or *35S: SpHERK1-3-nLUC* in *N. benthamiana*. The chemiluminescence images were obtained by applying 0.5 mM luciferin at 36 hpi. The experiment was repeated at least three times with similar results. MBP-nLUC and cLUC-GST were used as negative control. **E)** Domain organization of Pto, SpHERK1-1, SpHERK1-2, and SpHERK1-3. The features for domains were obtained from NCBI Conserved Domain Database and Uniprot database, and were ploted using IBS2.0. TM: transmembrane domain. **F)** SpHERK1-2$^{CD}$ phosphorylates AvrPtoB at Ser335 *in vitro* kinase assay. Recombinant GST-LecRK-IX.2$^{CD}$, MBP-SpHERK1-2$^{CD}$, MBP-AvrPtoB and site-mutated variants AvrPtoB$^{S335A}$ and AvrPtoB$^{T450D}$ were subjected to kinase activity detection assay. Protein phosphorylation was identified by immunoblot using α-pSer/Thr. The phosphorylated AvrPtoB was detected α-pS335). GST-Pto was used as positive control. **G, H)** SpHERK1-1-HA and SpHERK1-3-HA phosphorylate AvrPtoB at Ser335 by semi-*in vitro* kinase assay. *35S:SpHERK1-1-HA* or *35S:SpHERK1-3-HA* and *35S:GFP-HA* were transiently expressed in *N. benthamiana* leaves, respectively. The SpHERK1-1-HA, SpHERK1-3-HA and GFP-HA proteins were purified by anti-HA beads at 36 hpi, and were subjected to kinase activity detection along with MBP-AvrPtoB purified from *E. coli*. The phosphorylated AvrPtoB was detected by α-pS335.

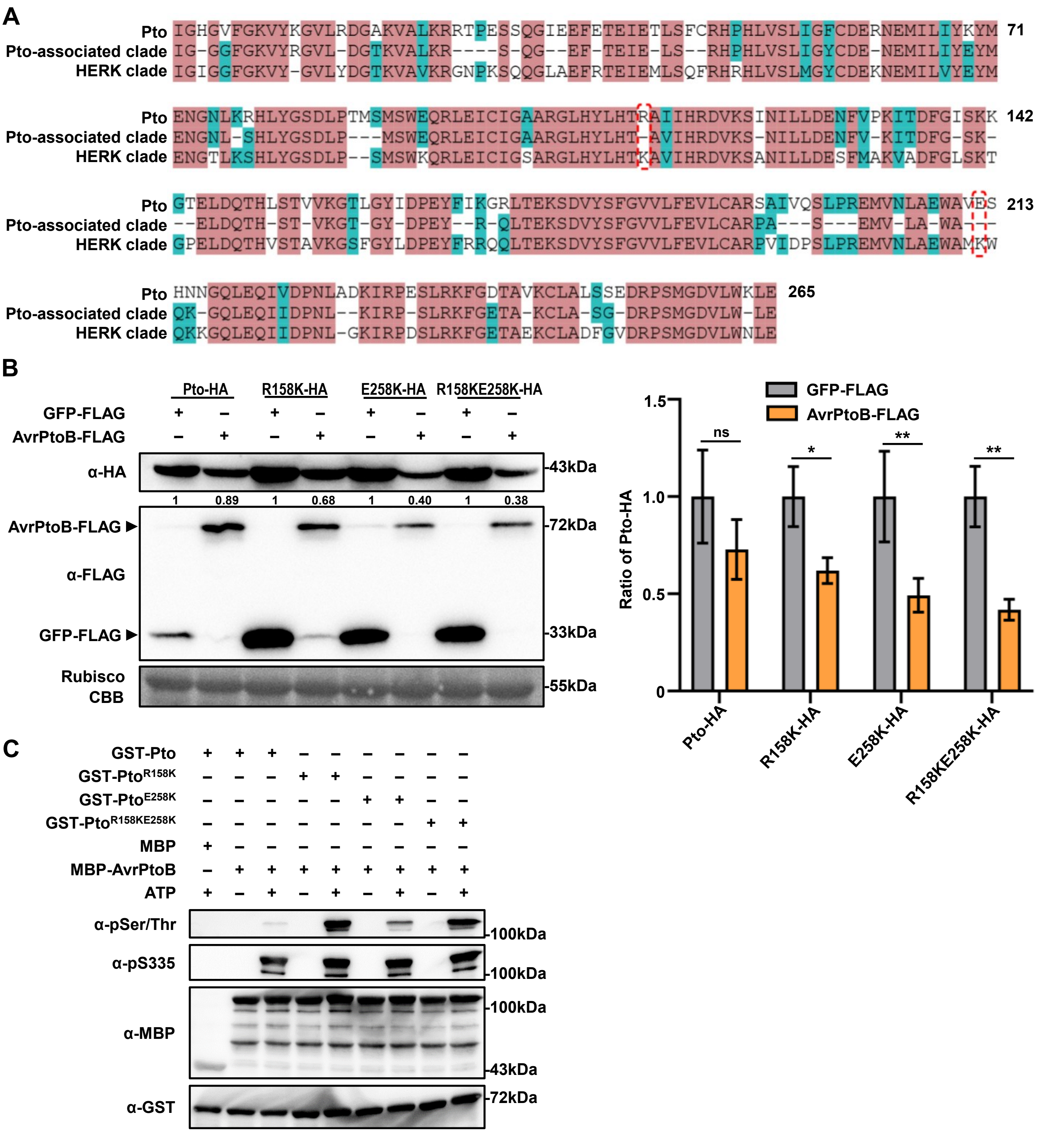


**Supplementary Figure S10. The amino acid sites Arg$^{158}$ and Glu$^{258}$ are important for the stability of Pto (Supports figure 7). A)** Alignment of 50% conserved sequences. The sequences alignment was performed using Pto kinase domain sequence, the 50% conserved sequence of all proteins in the Pto-associated clade to which Pto belongs, and the 50% conserved sequence of all proteins in the HERK clade. A dash ("-") indicated the absence of an amino acid or its conservation. The red dashed box highlights the site of interest where lysine was mutated to a non-lysine residue. Red amino acid backgrounds indicated positions where the sequences of "Pto," "Pto-associated clade," and "HERK clade" showed 100% identity, blue backgrounds represented positions with 60% identity, and white backgrounds indicated positions with 0% identity. **B)** The site mutation of Pto promotes it's degradation by AvrPtoB in *N. benthamiana*. *35S: Pto-HA*, *35S: Pto$^{R158K}$-HA*, *35S: Pto$^{E258K}$-HA*, *35S: Pto$^{R158K\ E258K}$-HA* was transiently expressed with *35S: AvrPtoB-FLAG* or *35S: GFP-FLAG*. Samples were collected after 36 hpi. The proteins were detected by immunoblotting using respective antibodies. CBB-stained Rubisco was used as loading control. Ratio of Pto-HA influenced by GFP-FLAG or AvrPtoB-FLAG was quantified by Image J. Values were means ± SD (n = 3 biological replicates; Two-way ANOVA; ns, no significance, *$p < 0.05$, **$p < 0.01$). **C)** The site mutation of Pto enhances it's phosphorylation capacity at AvrPtoB. GST-Pto, GST-Pto$^{R158K}$, GST-Pto$^{E258K}$, GST-Pto$^{R158KE258K}$, and MBP-AvrPtoB were subjected to *in vitro* kinase assay. Phosphorylation of AvrPtoB was detected by immunoblot using α-pSer/Thr and α-pS335 antibody.
